# Validated LC-MS/MS quantification of kynurenine pathway metabolites and pathway-balance ratios in brain tissue and microglial culture supernatant: application to neonatal stroke

**DOI:** 10.64898/2026.10.29.762714

**Authors:** Md Munnaf Hossen, Nee Edirisinghe, Emily Ross-Munro, Victor Mondal, Ritu Kumari, Abdullah Al Mamun Sohag, Isabelle K Shearer, Bobbi Fleiss, Rosita Zakaria

## Abstract

Kynurenine pathway (KP) metabolite imbalance is implicated in perinatal brain injury, and KP metabolites are measured in human plasma and cerebrospinal fluid as markers of neuroinflammation. Methods validated in neonatal brain tissue or glial culture supernatant to the same standard are scarce, limiting comparisons across preclinical and clinical measurements. We optimised LC-MS/MS detection of tryptophan (TRYP), kynurenine (KYN), kynurenic acid (KYNA), 3-hydroxykynurenine (3-HK), 3-hydroxyanthranilic acid (3-HAA), quinolinic acid (QA), anthranilic acid and picolinic acid after protein precipitation and Enhanced Matrix Removal-Lipid clean-up, validated it against Clinical and Laboratory Standards Institute (CLSI) acceptance criteria in brain tissue homogenate and cell culture supernatant, and applied it to a rat model of neonatal stroke and immune-activated BV2 microglia, reporting KYN/TRYP, KYNA/KYN, 3-HK/3-HAA and QA/KYNA ratios. The lower limit of quantification was 6.53-34.17 nM; imprecision was 0.57-11.47%, accuracy was 84.9-108.3%, recovery was 83.2-112.3%, and matrix effect was 87.8-108.8%, with linear calibration (R^2^ > 0.99) over 6.5-2000 nM. KYN/TRYP and QA/KYNA ratios were higher in the stroke hemisphere than in control brain at 1, 7 and 14 days (p<0.001), and KYNA/KYN was lower; lipopolysaccharide- and cytokine-stimulated BV2 microglia showed the same change in all four ratios (p<0.01). The assay quantifies eight KP metabolites and four pathway-balance ratios in brain tissue and microglial supernatant within clinical acceptance limits, allowing direct comparison with KP ratios reported in human plasma and cerebrospinal fluid. Changes in these ratios after neonatal stroke identify KYN/TRYP and QA/KYNA as candidate targets for neonatal biofluid studies.

## Introduction

The kynurenine pathway (KP) breaks down the essential amino acid tryptophan (TRYP) into bioactive metabolites that play vital roles in key biological processes, including oxidative stress, immune responses, and neurotransmission ^1, 2^. Specifically, the metabolites 3-hydroxykynurenine (3-HK) and 3-hydroxyanthranilic acid (3-HAA) can promote oxidative stress by generating reactive oxygen species, while quinolinic acid (QA) acts as an N-methyl-D-aspartate (NMDA) receptor agonist, potentially leading to excitotoxicity ^3, 4^. Conversely, the metabolite kynurenic acid (KYNA) is an NMDA receptor and alpha7-nicotinic acetylcholine receptor (α7 nAChR) antagonist; the KP metabolite, anthranilic acid (AA), exhibits antioxidant effects; and picolinic acid (PA) is neuroprotective by chelating metal ions and reducing oxidative stress ^5, 6^. The enzymes tryptophan-2,3-dioxygenase (TDO), indoleamine-2,3-dioxygenase-1 (IDO-1), and IDO-2 initiate the conversion of TRYP to kynurenine (KYN) ^7, 8^. The enzymes kynurenine aminotransferase (KAT), kynurenine 3-monooxygenase (KMO), and 3-hydroxyanthranilic acid dioxygenase (3-HAAO) subsequently act to convert KYN into KYNA, 3-HK and 3-HAA, and QA, respectively ^9, 10^.

Quantitative analysis of neuroactive KP metabolites is inherently complex due to their ultralow concentrations, diverse structural characteristics, chemical instability, and the presence of interfering substances in complex biological matrices, which can affect detection and accuracy ^11^. These difficulties are further exacerbated when analysing multiple metabolites or compounds, as each compound possesses distinct chemical properties and concentration levels ^12^. The combination of high-resolution chromatographic separation with highly sensitive and specific mass spectrometry has made liquid chromatography-mass spectrometry (LC-MS) a powerful tool for accurate metabolite quantification, overcoming challenges such as low concentrations, structural diversity, and matrix effects, even in complex biological samples. Additionally, optimised ionisation techniques and tailored mobile phase compositions enhance detection efficiency, ensuring reliable and reproducible measurements for both research and clinical applications ^13^.

Perinatal brain injury (PBI) is an umbrella term for damage to the developing brain due to insults such as hypoxic, inflammatory, oxidative, and excitotoxic processes, including those occurring after neonatal stroke ^14, 15^. PBI causes death or lasting neurocognitive and motor deficits in millions of infants each year ^14, 16, 17^. Growing evidence suggests that an imbalance in KP metabolites, with higher levels of neurotoxic metabolites (QA, 3-HAA, 3-HK) and lower levels of neuroprotective metabolites (AA, PA), is associated with brain damage following PBI ^18^. Microglia and astrocytes, the chief mediators of injury after PBI, are major sources of KP metabolites ^19^, and previous studies have suggested that KP dysfunction may act as a driver of injury in the developing brain ^20, 21^. However, many of the original studies assessed only a limited number of metabolites, leaving the balance between the two functional arms of the KP pathway unresolved. Finally, kynurenine is among the key metabolites emerging in cord/early blood metabolomics approaches used to support early identification of birth asphyxia ^22^. However, the precise contribution of KP metabolites to PBI progression and long-term neurodevelopmental outcomes remains unclear and will require improved high-throughput analytical approaches to be better understood.

Developing robust analytical methods requires comprehensive, standardised validation to ensure reproducibility, accuracy, and reliable inter-laboratory transferability, especially across diverse biological sample matrices. However, current LC-MS-based methods for the multiplex analysis of KP metabolites often lack consistent validation or evaluation across different sample matrices ^23^. These methodological weaknesses challenge inter-study comparisons and undermine confidence in reported metabolite concentrations, particularly for complex sample types such as brain tissue. Overcoming these limitations is critical for progress in KP research and for reliably measuring and interpreting metabolic alterations in both health and disease contexts. Consequently, this study aimed to develop and validate a reliable and transferable liquid chromatography-tandem mass spectrometry (LC-MS/MS) method for the integrated quantification of eight KP metabolites and pathway-balance ratios KYN/TRYP, KYNA/KYN, QA/KYNA, and 3-HK/3-HAA in two biological matrices with distinct interference profiles: brain tissue homogenates and cell culture supernatants. We then applied the method to brain tissue from a rat model of neonatal stroke and cell culture supernatants from immune-activated BV2 microglia to demonstrate its applicability across brain injury-relevant experimental models. We hypothesised that applying this fit-for-purpose method would detect coordinated changes in KP metabolites and pathway-balance ratios, reflecting a shift from neuroprotective to neurotoxic KP metabolism across both experimental paradigms.

## Materials and methods

### Chemicals

Pure compounds and standard materials including L-Tryptophan (Cat# T947210), rac-kynurenine (Cat# K661000), kynurenic acid (Cat# K660500), 2, 3-pyridinedicarboxylic acid (quinolinic acid, Cat# P991630), 3-hydroxy-DL-kynurenine (Cat# H950758), 3-hydroxyanthranilic acid (Cat# H801790), anthranilic acid (Cat# ANT-1077-10), picolinic acid (Cat# P437220), and internal standards for kynurenic acid-d3 (KYNA-d3, Cat# K660502) and 2, 3-pyridinedicarboxylic acid-d3 (QA-d3, Cat# P991633) were purchased from Toronto Research Chemicals, Toronto, ON, Canada. Other chemicals and solvents (LC-MS grade), such as formic acid (FA, Cat# A117-50), methanol (Cat# 047192.K7), acetonitrile (ACN) (Cat# 047138.K7), and isopropanol (Cat# 383920025), were purchased from Thermo Scientific (Waltham, MA, USA).

### Equipment

We used the Agilent ultra-high-performance liquid chromatography 1290-triple quadrupole mass spectrometry 6490 (UHPLC1290-QQQ6490) LC-MS/MS system (Agilent Technologies, Santa Clara, CA, USA; RRID:SCR_012482). For sample clean-up and chromatographic separation, we used Captiva EMR-Lipid (Agilent Cat# 5190-1002) and Poroshell 120 and EC-C18 (2.1×100 mm, 2.7 µm) columns (Agilent Cat# 695775-902), respectively. Extracted samples were dried using the Rotary Vacuum Centrifuge system (RVC2-25 CDplus; Martin Christ, Osterode am Harz, Germany) and centrifuged in a model 5427 R (Eppendorf SE, Hamburg, Germany).

### Preparation of standard materials

Based on the unique properties, including solubility and stability, solutions were prepared for TRYP, KYN, KYNA, QA, 3-HK, 3-HAA, AA, PA, KYNA-d3, and QA-d3 (**Supplementary Figure S1** & **Supplementary Table S1**) and stored at −80 °C for long-term use (6 months to 1 year). We selected two internal standards based on their structural similarity and comparable chromatographic and ionisation properties to the target analytes: KYNA-d3 quantified TRYP, KYN, KYNA, 3-HAA, AA, and PA, and QA-d3 quantified QA and 3-HK. Working stocks were prepared at 20,000 nM and stored at −20 °C for up to 2 weeks. For calibration standard preparation, working solutions were serially diluted with water to five concentrations ranging from 0 to 2000 nM.

### Preparation of samples and quality controls (QC)

Bottles and reusable glassware were washed with Milli-Q water and rinsed with methanol, dried in a 50 °C oven and finally cleaned with nitric acid solution (65%) followed by a Milli-Q water rinse before a final hot oven dry. For method validation, brain homogenates (15–25 mg starting material) from naive 10-day old Wistar rats (RRID:RGD_737929; see ethics details below) and Roswell Park Memorial Institute (RPMI) media containing 2 mM L-glutamine (Gibco, Thermo Fisher Scientific, Waltham, MA, USA; Cat# 11875) supplemented with 0.1% gentamicin (Thermo Fisher Scientific, 15750060) containing 10% of heat-inactivated newborn calf serum (NCS; Gibco, New Zealand, Lot: 2306210) (70 µL) were individually aliquoted into 1.5 mL Eppendorf tubes, and the final volume was made up to 100 µL/tube with distilled water. To each brain tissue quality control sample, we added 10 µL of a 50 nM ice-cold internal standard solution prepared in 100% methanol and 20 µL of KP standard solution (100–2000 nM) before extraction to minimise analyte degradation and ensure accurate quantification. The mixture was vortexed for 30 s, then 300 µL of ice-cold ACN was added and vortexed again for 20–30 s. To enhance protein precipitation efficiency, the samples were placed in a freezer at −20 °C for 10 min. Samples were then centrifuged at 10,000×*g* for 10 min at 4 °C, and approximately 350 µL of the clear supernatant was carefully transferred to a fresh tube, ensuring the protein pellet remained intact. Then, 200 µL of crash solvent (a 3:1 mixture of ACN and water with 1% FA) was added to the collected supernatant. The mixture was vortexed for 2–3 s and then passed through an Agilent Captiva EMR-Lipid cartridge, operated under a vacuum of 2–5 Hg, to ensure a controlled flow rate of one drop every 3–5 s. After elution, the vacuum pressure was increased for 1–2 min until the sample flow visibly slowed or ceased, ensuring maximum sample recovery.

Solvents were evaporated from the eluates using a rotary vacuum centrifuge at 37 °C and 5000 rpm, and the dried residues were reconstituted in 200 µL of mobile phase (0.1% FA in 95% water and 5% methanol) and vortexed for 30 s. Finally, 5 µL of the sample was injected into the LC-MS/MS system.

As external quality control materials for KP metabolites were not commercially available, in-house QC samples were prepared using filtered NCS. These QC samples were prepared and analysed separately alongside the cell culture supernatant samples during method validation to independently assess the accuracy of the analytical method. To prepare QC sample, 70 µL of filtered NCS was spiked with 20 µL of KP standards (100–2000 nM) and 10 µL of KYNA-d3 (50 nM) and processed using the same protocol as cell culture supernatant and brain samples. Similarly, neonatal stroke brain tissue (10–20 mg) and immune-stimulated BV2 culture medium (70 µL) were each spiked with 10 µL of 50 nM KYNA-d3 or QA-d3 and prepared as described above.

### LC-MS/MS analysis

LC-MS/MS analysis was conducted using an Agilent 1290 UHPLC and a 6490 QQQ MS/MS (RRID:SCR_012482). **Supplementary Tables S2 and S3** summarise the detailed specifications for the LC-MS/MS system. Chromatographic separation was achieved by a Poroshell 120 EC-C18 (2.1×100 mm, 2.7 µm) column at 38 °C. The mobile phase consisted of water containing 0.1% FA (A) and methanol containing 0.1% FA (B) at a flow rate of 0.3 mL/min. The mobile phase gradient elution was programmed as follows gradient-1: 0–5 min, 20% B; 5–6 min, 20–100% B; 6–8 min; 100% B; 8–9 min; 100–20% B and 9–12 min, 20% B for TRYP, KYN, KYNA, 3-HAA, PA, and AA, and gradient-2: 0–3 min, 5% B; 3–7 min, 5–40% B; 7–8 min; 40–100% B; 8–10 min; 100% B; and 10–13 min, 5% B for QA and 3-HK. We performed mass spectrometric detection using multiple reaction monitoring (MRM) with an electrospray ionisation (ESI) source in positive mode. The ionisation conditions were ion spray voltage 4500 V, temperature 250 °C, curtain gas 30 psi, nebuliser gas 20 psi, cell accelerator voltage 5.0 V, and sheath gas flow 11.0 L/min. Data acquisition and processing were performed using the Agilent MassHunter software package (version B.05.00/B5.0.291.0).

### Method validation

Method validation was conducted in accordance with the Clinical and Laboratory Standards Institute (CLSI) C62 guideline for LC-MS methods, which included assessments of linearity, imprecision, accuracy, recovery, matrix effects, and stability to ensure analytical reliability ^24–26^. We prepared each sample in triplicate at all QC levels and detected analytes using the validated LC-MS/MS method. For each targeted analyte, we used an internal standard to minimise matrix interference and ensure precise quantification, and we further optimised chromatographic or sample preparation conditions when interference was detected.

#### Linearity

Calibration and linearity were evaluated separately. QC quantification used five-point calibration curves spanning 0–2000 nM for TRYP, KYN, KYNA, 3-HAA, AA, and PA, and 0–500 nM for QA and 3-HK, while linearity was assessed independently using predefined criteria. We spiked matrix-matched filtered cell culture media at 10 and 2000 nM, and prepared concentrations below 10 nM by serial dilution of the 10 nM standard. We assessed nine concentrations (0.5, 10.5, 50.75, 105.0, 200.5, 420, 750, 1000, and 2000 nM). We included concentrations below the established lower limit of quantification (LLOQ) to characterise low-concentration response but excluded them from the validated quantitative range, which was defined by the analyte-specific LLOQ. Each concentration was analysed in triplicate within a single run across six independent days (n=6), and linearity was evaluated using LinChecker v1.1.

#### Sensitivity

Sensitivity was assessed by determining the lower limit of detection (LLOD) and LLOQ for each KP metabolite using the calibration-curve approach. We prepared and analysed a series of standard solutions ranging from 0.5 to 100 nM (n=20) to calculate the LLOD and LLOQ. We calculated the LLOD and LLOQ as 3.3σ/S and 10σ/S, respectively, where σ is the standard deviation of the analytical response and S is the slope of the calibration curve. The LLOQ was additionally required to demonstrate acceptable analytical performance, with imprecision within 20% of the nominal concentration ^27^.

#### Imprecision and accuracy

Intraday and interday imprecision and accuracy were assessed using cell culture supernatant and brain homogenate as sample-matched matrices, spiked with known concentrations of analytes same as three QC levels. For TRYP, KYN, KYNA, 3-HAA, AA, and PA, the QC levels were low QC (100 nM), medium QC (1000 nM; midpoint of the calibration curve), and high QC (2000 nM; close to the upper limit of the calibration range). For QA and 3-HK, the QC levels were low QC (100 nM), medium QC (250 nM), and high QC (500 nM). We tested each concentration level with six replicates. Intra- and inter-day imprecision were assessed using six replicates at three QC levels (low, medium, and high) within the same day and across six consecutive validation days ^28^. QC samples were analysed against a calibration curve, and the measured concentrations were compared with the nominal values to calculate accuracy and evaluate assay performance. Intraday and interday imprecision were determined as the coefficient of variation (CV) at each QC level, acceptable limits of ≤15% for medium and high QC levels and ≤20% for low QC levels ^29^.

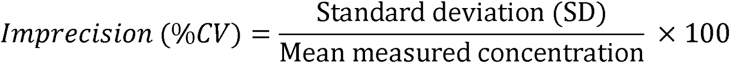

Accuracy was expressed as the percentage of the nominal value, with acceptable criteria set between 80–115% for medium and high QC levels and 70–120% for the low QC level ^30^.

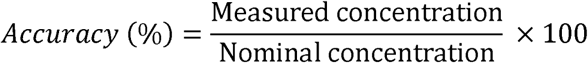

#### Matrix effect and recovery

We evaluated recovery and matrix effects to ensure the analytical method’s accuracy and robustness in complex biological matrices. Three QC concentration levels for cell culture supernatant and brain homogenate (low, medium, and high - outlined in the imprecision and accuracy section) were selected for analysis. For matrix-effect assessment, we used unspiked brain homogenate and cell culture medium as blank matrices to account for endogenous KP metabolites. We treated the endogenous analyte response in each unspiked matrix as the baseline and subtracted it from the corresponding response in the spiked matrix samples before calculating the matrix effect. We then created samples that were spiked with known concentrations of analytes (500 nM) before (pre-extraction spike) and after (post-extraction spike) the extraction process. To evaluate recovery, we spiked brain homogenate and cell culture media matrix samples with analytes before extraction (pre-extraction spike) and compared the results to the spiked standard concentration.

The matrix effect was calculated by comparing the concentration of analytes in post-extraction spiked samples to those in neat solutions using the equation ^31^:

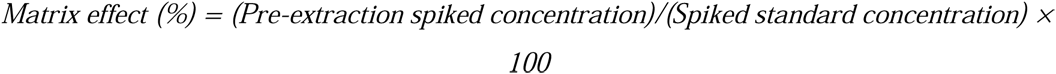

Values greater than 100% indicated ion enhancement, while values below 100% suggested ion suppression.

Recovery was determined by comparing the peak areas of analytes in pre-extraction spiked samples to post-extraction spiked samples using the formula ^31^:

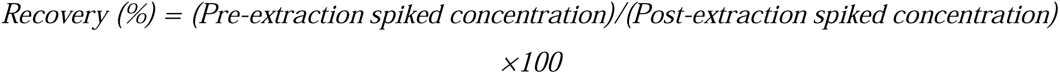

Acceptable recovery was between 80–115%. Matrix effects were deemed acceptable if the deviation was within ±15%. Results confirmed that the method maintained high reproducibility and minimal matrix interference.

#### Stability

Stability analysis evaluated the effects of a single round of sample freezing and thawing, followed by short- and long-term storage. Samples were assessed following storage at room temperature for up to 24 h and at −20 °C for up to 21 days. Stability testing used KP standard spiked concentrations of 2000 nM for TRYP and 100 nM for KYN, KYNA, QA, 3-HK, 3-HAA, PA, and AA, including the internal standard. We compared results with freshly prepared samples and considered analytes stable when accuracy remained within ±15% ^32, 33^.

### Animal model and sample collection

We conducted all experimental procedures with prior approval from the RMIT University Animal Ethics Committee (AEC 27678) and reported them in accordance with the ARRIVE guidelines. We obtained time-mated Wistar rats from Australian BioResources (Perth, Western Australia) at 12 to 16 days of pregnancy and allowed them to deliver undisturbed. We recorded body weights for all pups on postnatal day 8, assigned each pup a random identifier, and allocated pups to groups within litters in a block design using restricted randomisation. We applied our validated photothrombotic protocol to induce a focal ischemic lesion that replicates facets of neonatal stroke in Wistar rats (RRID:RGD_737929) ^34^. Briefly, at postnatal day 10 (P10), pups were anaesthetised with 5% isoflurane (Isoflo; Abbott, NSW, Australia) in 0.4 L/min oxygen by face mask for 2 min, with anaesthesia maintained at 2 to 3% isoflurane in 0.4 L/min oxygen, and Rose Bengal (25 mg/kg, intraperitoneally; Merck, Australia; Cat# 330000) was administered. Five minutes after administration, photothrombotic injury was induced by illuminating the skin overlying the motor cortex for 10 min with a high-intensity 565 nm light-emitting diode (LED; Thorlabs, Newton, NJ, USA; Cat# M565F3) focused to a 1 mm diameter area and secured in position using an arm attached to the stereotaxic frame. Control animals received Rose Bengal but were not exposed to LED illumination. Brain tissue samples of a fixed size were collected at post-stroke 24 h, 7 days, and 14 days using a 3.0 mm diameter biopsy punch centred on the lesion in the ipsilateral hemisphere and a comparable region of motor cortex in the contralateral hemisphere (non-lesioned side) (**Fig. 1**) or from the motor cortex of a control animal (no injury). We immediately snap-froze samples in liquid nitrogen and stored them at −80 °C until analysis. We culled four male and four female pups (n=8 pups) from four litters per group at each time point. We used four pups (two males and two females) per group for LC-MS/MS analysis, and the remaining four pups (two males and two females) for neuropathological analysis. Sex does not influence the injury profile in this model ^34^. As this study aimed to validate the utility of this method, the sample size was determined based on previous experience with this model in which injury is substantial and consistent. Although the researcher handling the tissue was blinded to the specific group allocation, with labelling using only litter and pup numbers, the lesion was highly visible in all analyses. At tissue collection, pups were anaesthetised with an intraperitoneal injection of Lethabarb (325 mg/mL pentobarbitone sodium, >200 mg/kg; Virbac, Australia) with lignocaine (10 mg/mL; Cenvet, Australia). Pups for LC-MS/MS were decapitated, and brain tissue collected fresh; pups for neuropathology were transcardially perfused with 0.1 M phosphate-buffered saline (PBS, pH 7.4) for 1 min followed by 4% paraformaldehyde (PFA; Scharlau, Barcelona, Spain) and processed after 24 hours in 4% paraformaldehyde to paraffin.

**Fig. 1:**
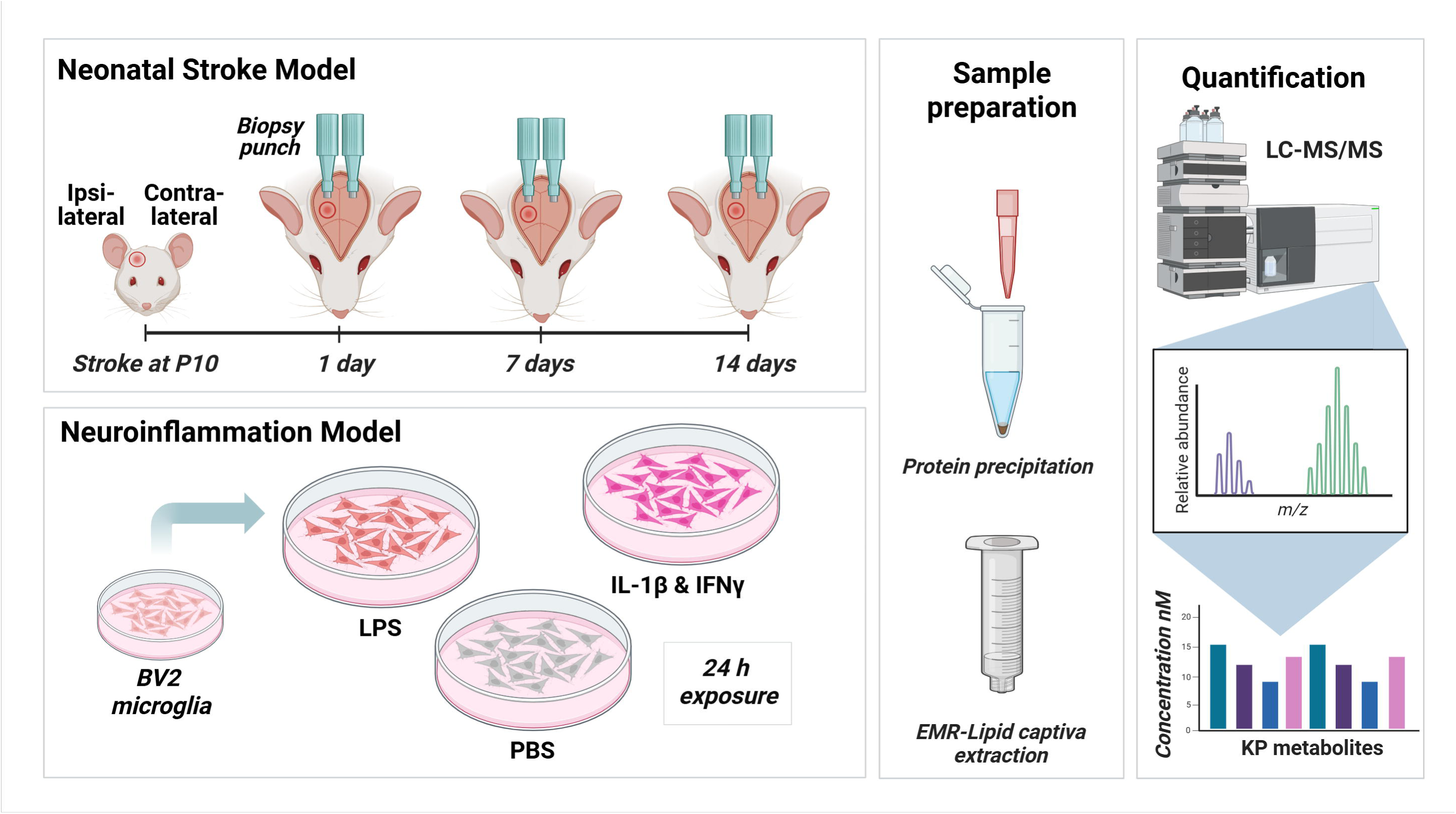
Schematic illustration of sample collection, preparation, and LC–MS/MS analysis of kynurenine pathway metabolites from the in vivo neonatal stroke rat model and the in vitro BV2 microglial neuroinflammation model.

### Haematoxylin and eosin (H&E) staining and lesion volume analysis

We collected, fixed and paraffin-embedded brains and prepared them as in our previous studies ^34^. We deparaffinised and rehydrated every twentieth coronal section (8 μm) spanning the lesion, then stained it with 0.3% Mayer’s haematoxylin (Amber Scientific, Australia) for 4 min. Slides were rinsed thoroughly under running tap water, immersed in Scott’s tap water substitute (20 g magnesium sulphate and 2 g sodium bicarbonate dissolved in 1 L deionised water; Sigma-Aldrich, St. Louis, MO, USA) for 1 min, then washed again in tap water for 3 min. Sections were counterstained with 1% alcohol eosin (Amber Scientific, Australia) for 3 min, dehydrated through graded alcohols, and mounted with dibutylphthalate xylene (DPX; Merck, Australia). Brightfield images were acquired at 10× magnification using an Olympus VS120 Virtual Slide Microscope (Olympus Life Science, VIC, Australia, RRID:SCR_018411). We quantified stroke lesion volume using ImageJ software (Fiji, version 2.0; NIH, Bethesda, MD, USA, RRID: SCR_002285) following an established laboratory protocol ^34^.

### Cell culture

BV2 (RRID: CVCL_0182), an immortalised mouse microglial cell line, was sourced from Elabscience (Wuhan, China; Cat# EP-CL-0493). Cells were cultured following our laboratory’s previously described method ^35, 36^ in RPMI media containing 2 mM L-glutamine (Gibco, Cat# 11875) supplemented with 0.1% gentamicin (Thermo Fisher Scientific, Cat# 15750060) and 10% heat-inactivated NCS. After detachment with TrypLE Express (Gibco, Cat# 12604021), cells were plated at 20,000 cells/cm^2^ in 48-well plates in 200 µL of RPMI media (Gibco, Cat# 11875). All experiments used BV2 cells from passage numbers 4–6. Twelve hours after plating, BV2 cells were treated with inflammatory agents: lipopolysaccharide (LPS) (1 µg/mL; O55:B5 from *Escherichia coli*; Sigma-Aldrich, Cat# L6529), or an inflammatory cytokine mix (ICM), IL-1β (50 ng/mL; Miltenyi Biotec, Bergisch Gladbach, Germany; Cat# 130-101-682) and IFN-γ (20 ng/mL; Miltenyi Biotec, Cat# 130-105-785), or a PBS-only vehicle (**Fig. 1**). After 24 h of treatment, the supernatant was collected and stored at −80 °C for LC-MS/MS analysis and cells were collected for RNA extraction as detailed below.

### Gene expression studies on BV2-microglia

We performed RNA extraction, cDNA synthesis, and qRT-PCR according to established laboratory protocols ^37, 38^. In brief, total RNA was extracted using the Qiagen kit (Qiagen, Hilden, Germany; Cat# 74136) in 350 µL of RLT lysis buffer (Qiagen, Cat# 74136), which was split across three replicate wells and stored at −80 °C until RNA was extracted. RNA had an A260/280 ratio of 1.8–2.1 and an A260/230 ratio >1.8, as determined with a NanoDrop One Microvolume UV-Vis Spectrophotometer (Thermo Fisher Scientific; RRID:SCR_023005).

For cDNA synthesis, 1 μg of RNA was reverse transcribed into cDNA using the iScript cDNA Synthesis Kit (Bio-Rad, Hercules, CA, USA; Cat# 1708891) with the Bio-Rad T100 instrument (RRID:SCR_021921). We designed primers using Primer3Plus software (RRID:SCR_003081) and Primer-BLAST (RRID:SCR_003095). Primers (Thermo Fisher Scientific, Australia) were validated to have an efficiency range of 80% to 120%. Primer sequences can be found in **Table 1**.

**Table 1:**
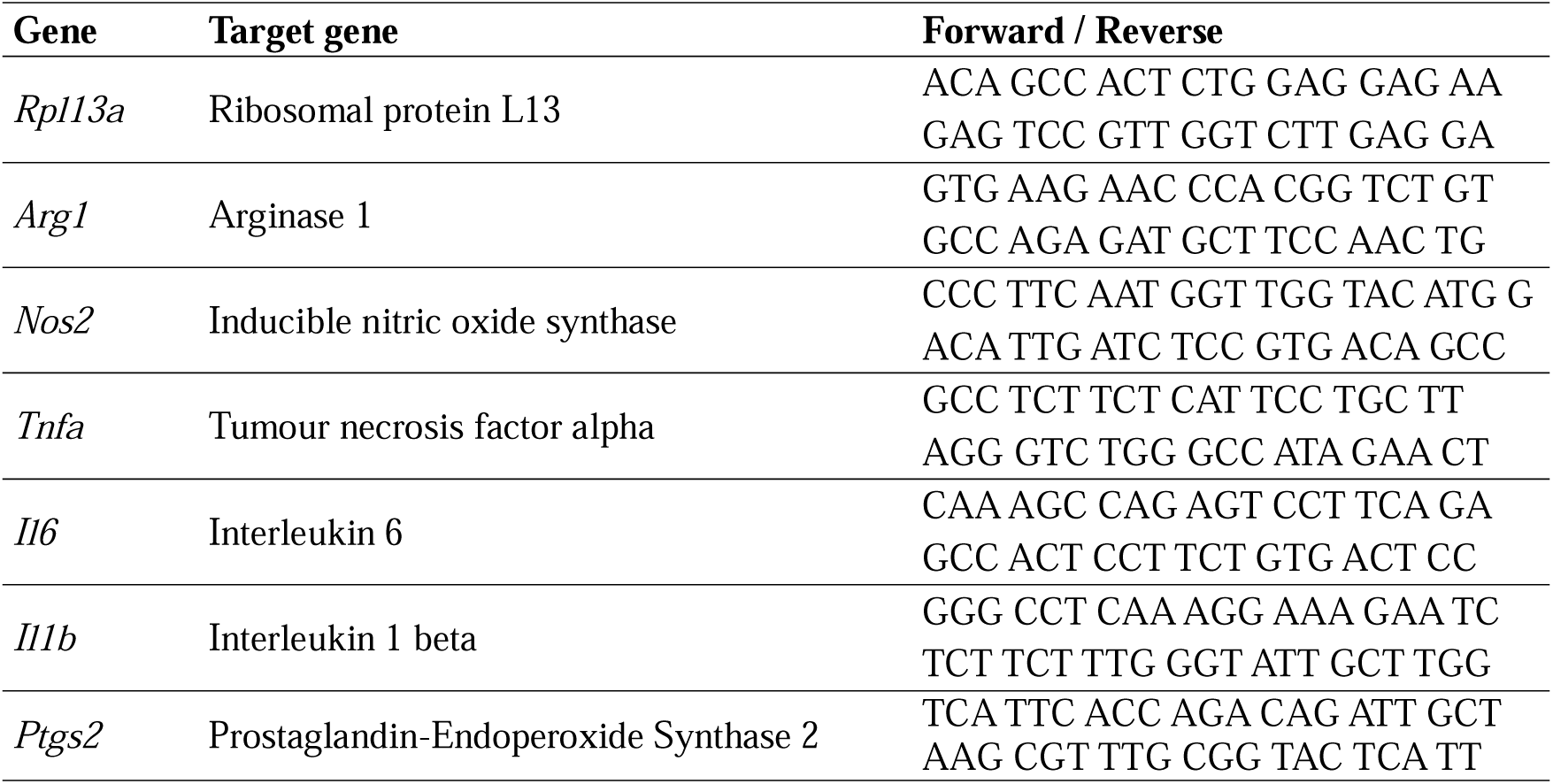
List of primer sequences used for inflammatory and anti-inflammatory gene expression analysis.

For qRT-PCR, we used iTaq Universal SYBR Green Supermix (Bio-Rad, Cat# 1725124) with 1 µL cDNA to a final volume of 20 µL in a 384-well plate, with triplicate reactions. The qRT-PCR was performed on the QuantStudio 7 Flex instrument (Applied Biosystems, Foster City, CA, USA; RRID:SCR_020245) using a 40-cycle two-step program with 5 s denaturation at 95 °C and 10 s annealing at 60 °C. *Rpl13a* was used as the reference gene based on our previous studies ^38^. mRNA expression was calculated using the 2^−ΔΔCt^ method.

### Data analysis and statistics

Data acquisition, visualisation, and analysis were performed using Agilent MassHunter software (version B.05.00/B5.0.291.0; RRID:SCR_015040). Linearity was evaluated using LinChecker software (version 1.1). Data from neonatal stroke brain tissue and BV2 cell culture supernatants are presented as mean ± SEM, and statistical analyses were conducted using GraphPad Prism (version 10.1.0; RRID:SCR_002798). To assess injury effects across hemispheres (control, contralateral, and ipsilateral) and post-stroke time points, we applied a mixed-effects analysis followed by Sidak’s *post hoc* test for multiple comparisons to neonatal brain tissue data. For infarct volume data and BV2 cell culture experiments, we used one-way ANOVA followed by Sidak’s *post hoc* test to assess infarct volume and evaluate the effects of inflammatory stimulation across treatment groups (PBS, LPS, and ICM). Statistical significance was defined as p<0.05. No samples from the animal or cell culture experiments were excluded from the analyses for any reason. We checked normality and homogeneity of variance of the residuals before the mixed-effects analysis and ANOVA.

## Results

### Optimisation and development of the LC-MS/MS method

#### Mass spectrometry analysis

We optimised the MRM transition and collision energies for each KP metabolite in both positive and negative ionisation modes. We selected two product ions for each metabolite (Table 2), designating the first, the most abundant and robust, as the quantifier and the second as the qualifier. All KP metabolites produced detectable precursor and product ions in both positive and negative ionisation modes. However, positive ionisation produced more consistent signal intensity and sensitivity across analytes. Therefore, we established the LC-MS/MS method in positive ionisation mode to ensure robust, uniform metabolite detection.

**Table 2:** Summary of MRM conditions of KP metabolites in positive ionisation mode.

| Metabolites | Precursor Ion<br>[M+H]<sup>+</sup> | 1st Product ion (quantifier) |  | 2nd Product ion (qualifier) |  |
| --- | --- | --- | --- | --- | --- |
|  |  | m/z | Collision energy (eV) | m/z | Collision energy (eV) |
| TRYP | 205 | 188 | 3 | 146 | 13 |
| KYN | 209 | 192 | 5 | 146 | 16 |
| KYNA | 190 | 144 | 19 | 116 | 34 |
| KYNA-d3 | 195 | 149 | 19 | 121 | 35 |
| QA | 168 | 150 | 5 | 106 | 13 |
| QA-d3 | 171 | 153 | 5 | 109 | 13 |
| 3-HK | 225 | 208 | 5 | 110 | 23 |
| PA | 124 | 106 | 10 | 78 | 21 |
| 3-HAA | 154 | 136 | 9 | 108 | 20 |
| AA | 138 | 120 | 10 | 92 | 21 |

**Abbreviations:** Tryptophan (TRYP), kynurenine (KYN), kynurenic acid (KYNA), kynurenic acid d3 (KYNA-d3), quinolinic acid (QA), 3-hydroxykynurenine (3-HK), 3-hydroxyanthranilic acid (3-HAA), anthranilic acid (AA), picolinic acid (PA), kynurenic acid (KYNA), and quinolinic acid-d3 (QA-d3).

#### Liquid chromatography separation

We optimised the mobile phase composition to improve sensitivity and chromatographic performance for each KP metabolite. A combination of 0.1% FA in water (mobile phase A) and 0.1% FA in methanol (mobile phase B) showed optimal peak intensity, reasonable resolution, shape, and elution times. Additionally, the water-methanol mobile-phase combination showed greater separation efficiency, yielding sharper peaks and improved resolution, making it the preferred choice for reliable and reproducible quantification of KP metabolites compared with the FA-water-acetonitrile mixture. Next, we developed and refined two mobile-phase gradients to improve KP metabolite separation and elution. With the finalised gradient 0–5 min, 20% B; 5–6 min, 20–100% B; 6–8 min; 100% B; 8–9 min; 100–20% B and 9–12 min, 20% B (gradient 1) for TRYP, KYN, KYNA, 3-HAA, PA, and AA, then, 0–3 min, 5% B; 3–7 min, 5–40% B; 7–8 min; 40–100% B; 8–10 min; 100% B; and 10–13 min, 5% B (gradient 2) was optimised for QA and 3-HK, and all eight KP metabolites and the internal standard were efficiently separated within 7 min, with a total runtime of 13 min per analysis (Supplementary Figure S2). Additionally, LC parameters were tested, including an injection volume of 5 µL, a 0.3 mL/min flow rate, and a column oven temperature of 38 °C.

### Method validation and analytical performance

#### Assay analytical range, calibration curve, linearity and sensitivity

We prepared calibration curves for targeted metabolites over a five-point concentration range: 0 to 2000 nM for TRYP, KYN, KYNA, 3-HAA, PA, and AA; and 0 to 500 nM for QA and 3-HK. Table 3 and Supplementary Figures S3 and S4 summarise the linear assessment ranges and linear regression coefficient (R^2^) values. The optimised method showed linear assessment ranges of 0.5–2000 nM (R^2^ > 0.99) for TRYP; 0.5–1000 nM for KYN, KYNA, 3-HAA, PA, and AA; and 0.5–500 nM for QA and 3-HK. In the brain matrix, the LLOD for KP metabolites ranged from 2.13 to 6.65 nM, and the LLOQ ranged from 6.53 to 19.90 nM. Similarly, the LLOD for the cell culture matrix ranged from 2.37 to 11.38 nM, and the LLOQ ranged from 7.11 to 34.17 nM (**Table 3**).

**Table 3:**
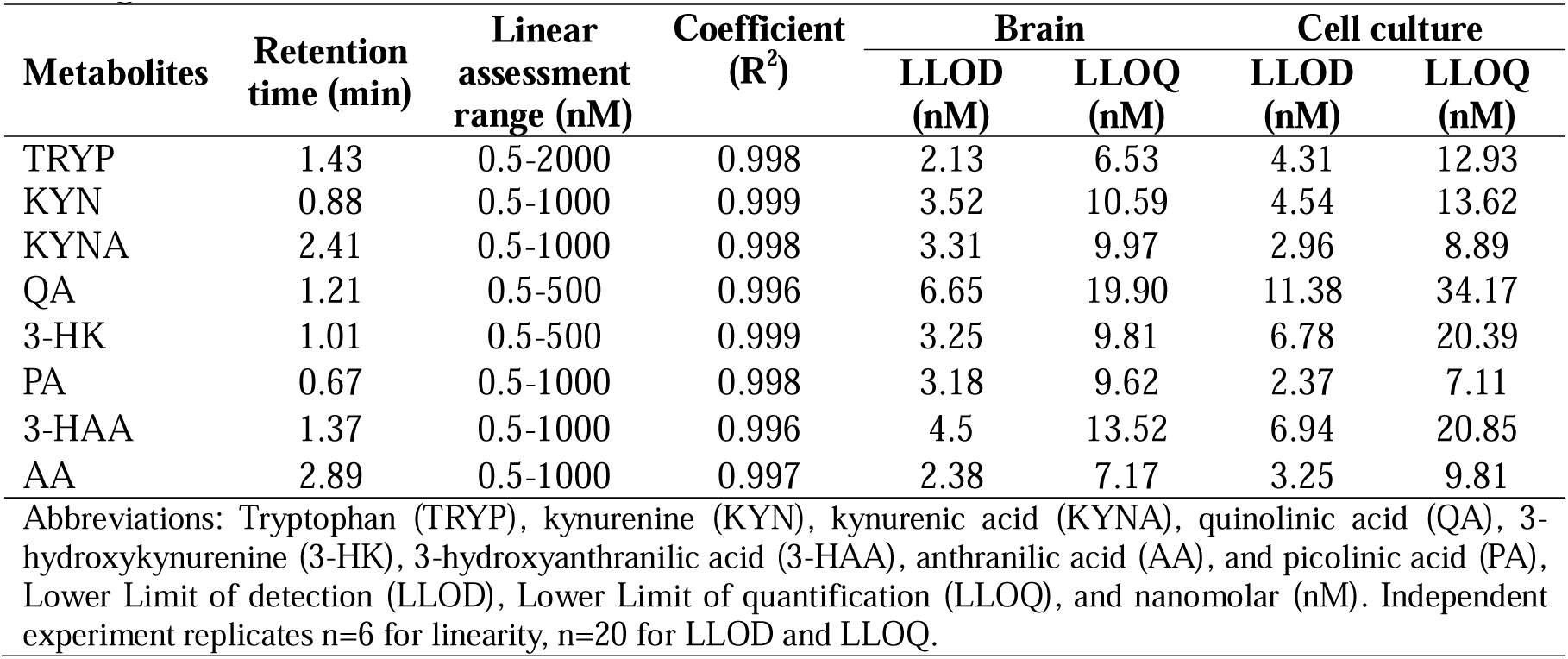
Summary of retention time, calibration range, and coefficient for both brain homogenate and cell culture media.

| Metabolites | Retention time (min) | Linear assessment range (nM) | Coefficient ($R^2$) | Brain | | Cell culture | |
| --- | --- | --- | --- | --- | --- | --- | --- |
|  |  |  |  | LLOD (nM) | LLOQ (nM) | LLOD (nM) | LLOQ (nM) |
| TRYP | 1.43 | 0.5-2000 | 0.998 | 2.13 | 6.53 | 4.31 | 12.93 |
| KYN | 0.88 | 0.5-1000 | 0.999 | 3.52 | 10.59 | 4.54 | 13.62 |
| KYNA | 2.41 | 0.5-1000 | 0.998 | 3.31 | 9.97 | 2.96 | 8.89 |
| QA | 1.21 | 0.5-500 | 0.996 | 6.65 | 19.90 | 11.38 | 34.17 |
| 3-HK | 1.01 | 0.5-500 | 0.999 | 3.25 | 9.81 | 6.78 | 20.39 |
| PA | 0.67 | 0.5-1000 | 0.998 | 3.18 | 9.62 | 2.37 | 7.11 |
| 3-HAA | 1.37 | 0.5-1000 | 0.996 | 4.5 | 13.52 | 6.94 | 20.85 |
| AA | 2.89 | 0.5-1000 | 0.997 | 2.38 | 7.17 | 3.25 | 9.81 |

Abbreviations: Tryptophan (TRYP), kynurenine (KYN), kynurenic acid (KYNA), quinolinic acid (QA), 3-hydroxykynurenine (3-HK), 3-hydroxyanthranilic acid (3-HAA), anthranilic acid (AA), and picolinic acid (PA), Lower Limit of detection (LLOD), Lower Limit of quantification (LLOQ), and nanomolar (nM). Independent experiment replicates n=6 for linearity, n=20 for LLOD and LLOQ.

#### Imprecision, accuracy, recovery, and matrix effect

We tested imprecision using matrix-matched cell culture and brain samples at three concentrations (low, medium, and high, as outlined in the method section), in both within-day (intraday) and between-day (interday) runs. For KP metabolites, intraday imprecision (CV%) for QC high-to-low concentrations ranged from 0.57% to 9.33%, and interday imprecision ranged from 0.92% to 11.47% in the brain matrix, as summarised in **Table 4**. Similarly, in cell culture media, intraday and interday imprecision for the high-to-low QC range (CV%) were 1.44–7.59% and 1.54–11.41%, respectively (**Table 5**).

**Table 4:** Summary of imprecision, accuracy, recovery, and matrix effect for the brain homogenate.

| Metabolite | Imprecision |  | Accuracy<br>(%±SD) | Recovery<br>(%±SD) | Matrix effect<br>(%±SD) |
| --- | --- | --- | --- | --- | --- |
|  | Intraday<br>(high to low,<br>CV%) | Interday (high<br>to low, CV%) |  |  |  |
| TRYP | 1.53 - 5.30 | 1.34 - 5.47 | 102.1±5.33 | 100.82±0.83 | 98.19±1.25 |
| KYN | 0.57 - 3.93 | 0.92 - 5.85 | 97.0±5.96 | 91.0±4.33 | 98.05±1.33 |
| KYNA | 1.66 - 5.16 | 1.82 - 5.21 | 102.9±4.81 | 103.1±5.57 | 89.65±2.82 |
| QA | 2.60 - 6.30 | 3.68 - 9.44 | 84.9±5.38 | 83.2±5.18 | 95.24±2.34 |
| 3-HK | 1.36 - 9.33 | 2.72 - 11.47 | 89.2±4.54 | 90.5±4.21 | 96.69±0.41 |
| PA | 0.58 - 6.62 | 1.13 - 7.37 | 101.4±6.72 | 96.7±2.70 | 89.49±1.73 |
| 3-HAA | 1.35 - 6.81 | 1.69 - 8.28 | 90.8±3.26 | 90.8±3.49 | 97.97±1.80 |
| AA | 1.42 - 7.54 | 1.66 - 9.14 | 108.3±3.91 | 96.8±3.71 | 95.08±0.56 |

**Abbreviations:** Tryptophan (TRYP), kynurenine (KYN), kynurenic acid (KYNA), Quinolinic acid (QA), 3-hydroxykynurenine (3-HK), 3-hydroxyanthranilic acid (3-HAA), anthranilic acid (AA), and picolinic acid (PA). Data presented as mean ± SD, independent experiment replicates n=6.

**Table 5:** Summary of imprecision, accuracy, recovery, and matrix effect, for cell culture media.

| Metabolite | Imprecision |  | Accuracy<br>(%±SD) | Recovery<br>(%±SD) | Matrix effect<br>(%±SD) |
| --- | --- | --- | --- | --- | --- |
|  | Intraday<br>(high to low,<br>CV%) | Interday<br>(high to low,<br>CV%) |  |  |  |
| TRYP | 1.64 - 5.51 | 1.88 - 6.49 | 94.58±2.44 | 102.49±8.41 | 108.78±10.12 |
| KYN | 1.89 - 5.67 | 2.13 - 5.47 | 95.65±1.92 | 88.13±1.26 | 96.79±1.59 |
| KYNA | 1.49 - 5.82 | 2.07 - 7.14 | 95.92±1.90 | 101.75±1.18 | 91.25±1.17 |
| QA | 1.52 - 6.76 | 1.42 - 11.41 | 90.84±1.83 | 89.45±0.75 | 87.88±0.81 |
| 3-HK | 3.22 - 7.59 | 4.42 - 8.54 | 91.53±3.05 | 97.97±0.52 | 92.68±3.59 |
| PA | 1.71 - 5.93 | 2.04 - 6.59 | 92.06±1.80 | 86.66±0.82 | 92.99±2.44 |
| 3-HAA | 1.44 - 5.26 | 1.77 - 7.72 | 92.48±1.89 | 96.01±0.18 | 95.57±1.27 |
| AA | 1.46 - 6.00 | 1.54 - 7.90 | 95.54±2.25 | 112.31±5.06 | 106.37±8.25 |

**Abbreviations:** Tryptophan (TRYP), kynurenine (KYN), kynurenic acid (KYNA), Quinolinic acid (QA), 3-hydroxykynurenine (3-HK), 3-hydroxyanthranilic acid (3-HAA), anthranilic acid (AA), and picolinic acid (PA). Data presented as mean ± SD, independent experiment replicates n=6.

We evaluated accuracy using samples from the imprecision experiments and determined it using the standard addition method. Tables 4 and 5 present the accuracy of KP metabolites, showing values ranging from 89.2% to 108.3% in the brain matrix and from 90.84% to 95.92% in the cell culture matrix. The results also demonstrated mean recovery rates of 83.2–103.1% in the brain matrix and 86.66–112.31% in the cell culture matrix.

Co-eluting compounds from brain and cell culture matrices affected target KP analyte ionisation differently, causing matrix effects. In the brain matrix, matrix effects ranged from 89.49% to 98.19%, with PA showing the greatest ion suppression and TRYP the least (**Table 4**). Similarly, in the cell culture matrix, effects ranged from 87.88% to 108.78%, with QA exhibiting the highest ion suppression and TRYP the least affected (**Table 5**).

#### Stability

Stability for all targeted KP metabolites was tested at room temperature and −20 °C, reported in **Fig. 2** as the percent change in concentration compared to the initial concentration. All eight KP metabolites are stable at room temperature for 24 h. After freezing and thawing, TRYP, KYN, KYNA, PA, and AA were stable at −20 °C for up to 21 days in both matrices. In contrast, at −20 °C, the concentrations of QA, 3-HK, and 3-HAA decreased by more than 15% after 14 days relative to day 1 levels. These findings support analysis of 3-HK, 3-HAA, and QA within 7 days when stored at −20 °C, while snap-freezing and storage at −80 °C are recommended for longer-term storage and analysis.

**Fig. 2:**
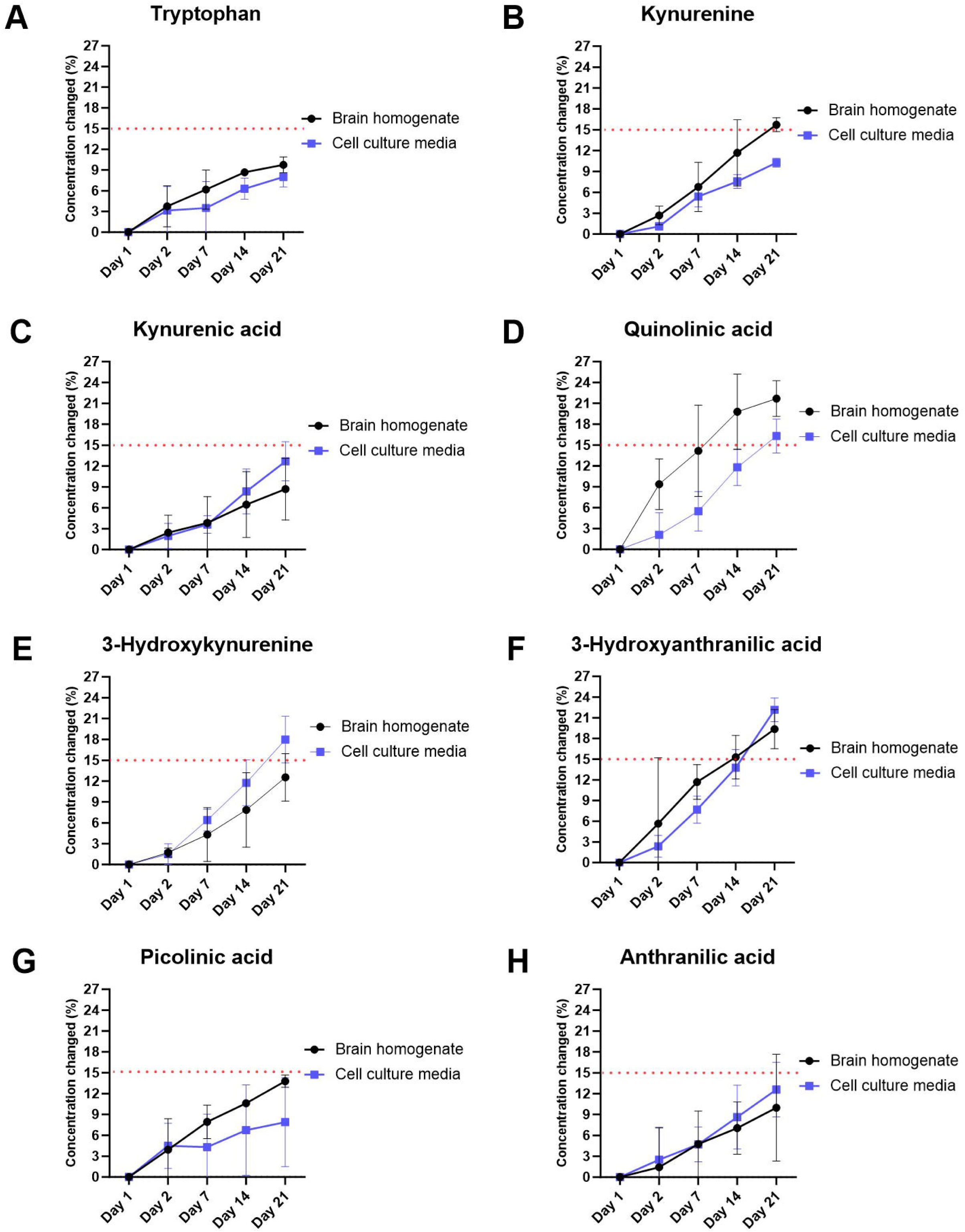
Stability analysis of KP metabolites in the brain and cell culture matrices at −20 °C up to 21 days. Concentration changes (%) of a) tryptophan, b) kynurenine, c) kynurenic acid, d) quinolinic acid, e) 3-hydroxykynurenine, f) 3-hydroxyanthranilic acid, g) picolinic acid, and h) anthranilic acid. Data presented as Mean ± standard error of the mean (SEM), independent replicates, n=3.

### Application of the validated method to biological samples

Ischemia and microglia-mediated inflammatory responses are well-recognised features of brain injury. To assess the practical applicability of our validated LC-MS/MS method, we quantified KP metabolites in an *in vivo* model of neonatal focal ischemic stroke and in an *in vitro* model of microglia-mediated inflammation. Pathway-balance ratios quantify four features of KP flux: entry into the KP (KYN/TRYP), diversion to the neuroprotective branch (KYNA/KYN), oxidative balance (3-HK/3-HAA), and diversion to the neurotoxic branch (QA/KYNA).

One-way ANOVA of the neuropathological data showed a significant main effect of time on infarct volume (p<0.001), with infarct size largest at 24 h (4.9±0.67 mm^3^) and smallest at 14 days post-injury (0.19±0.06 mm^3^) (**Fig. 3A).** We then applied the LC-MS/MS method to quantify KP metabolites in brain tissue collected at 24 h, 7 days, and 14 days after injury, and in age-matched uninjured controls. Mixed-effects analysis revealed a significant main effect of hemisphere source (p<0.001) and time post-stroke (p<0.05) on measured KP metabolite levels, except AA and PA (**Fig. 3B–D**).

**Fig. 3:**
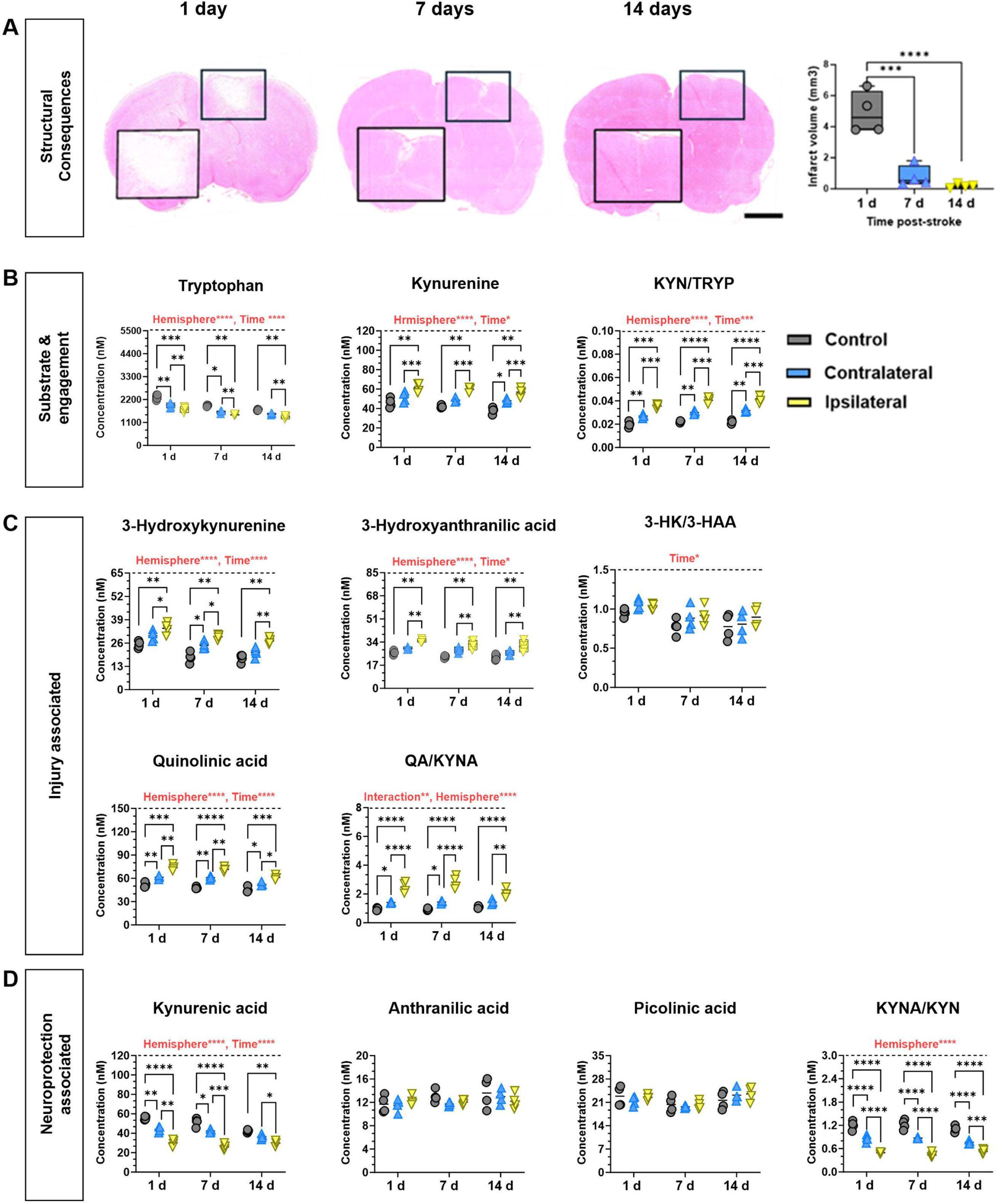
Lesion volume and kynurenine pathway metabolite alterations following neonatal focal ischemic injury. (A) Representative haematoxylin and eosin (H&E) stained coronal brain sections showing infarct location and quantification of infarct volume at 1 and 7 and 14 days after injury. (**B**) Concentrations of the KP metabolites associated with substrate and pathway engagement (tryptophan [TRYP], kynurenine [KYN] and the kynurenine/tryptophan ratio), in (**C**) metabolites associated with injury progression (3-hydroxykynurenine [3-HK], 3-hydroxyanthranilic acid [3-HAA], the 3-HK/3-HAA ratio, quinolinic acid [QA] and the QA/KYNA ratio) and in **D**) metabolites associated with a neuroprotective response (kynurenic acid [KYNA], anthranilic acid [AA], and picolinic acid [PA], and KYNA/KYN ratio) measured in brain tissue from control animals (uninjured) and from the contralateral (uninjured from stroke animal) and ipsilateral hemispheres (stroke hemisphere) at each time point. Data are presented as mean ± SEM; n = 4 litters; n = 4 pups per group, shown as individual points on the graphs; outlined in Supplementary Table S4, S5, and S6. Statistical significance was determined using a one-way ANOVA (A) and a mixed effect analysis followed by Šídák’s multiple comparisons test (B-D). Significance levels in the post-test are indicated on the graphs (*p < 0.05, **p < 0.01, ***p < 0.001, ****p < 0.0001).

In Sidak’s *post hoc* tests, TRYP (**Fig. 3B**) was markedly decreased in the ipsilateral (injured) hemisphere (p<0.01) and the contralateral hemisphere (p<0.05) at 1 and 7 days relative to control levels. At 1, 7, and 14 days after stroke induction, TRYP levels were significantly lower in the ipsilateral hemisphere compared with the contralateral hemisphere (p<0.01). In contrast, *post hoc* tests demonstrated that KYN levels were increased at all time points in the ipsilateral hemisphere compared with controls (p<0.01) **(Fig. 3B).** At 1, 7, and 14 days after stroke, KYN levels were significantly elevated in the ipsilateral hemisphere compared with the contralateral hemisphere (p<0.001). Additionally, at 14 days after injury, KYN levels in the contralateral hemisphere were higher than in controls (p<0.05).

In the Sidak’s *post hoc* tests, the neuroprotective metabolite KYNA (**Fig. 3D**) was lower in the ipsilateral hemisphere across all time points and was significantly reduced compared with both the contralateral hemisphere (p<0.05) and controls (p<0.01). KYNA levels in the contralateral hemisphere were also consistently lower than controls at 1 and 7 days post-stroke (p<0.05). For the neurotoxic metabolites 3-HK, 3-HAA, and QA, *post hoc* analysis revealed that levels were significantly higher in the ipsilateral hemisphere compared with controls across all time points (p<0.01) (**Fig. 3C**). At 1 and 7 days post-stroke, 3-HK and QA levels were significantly elevated in the ipsilateral hemisphere compared with the contralateral hemisphere (p<0.05). Additionally, QA levels in the contralateral hemisphere were higher than controls at 1, 7, and 14 days post-stroke (p<0.05).

Mixed-effects analysis revealed a significant hemisphere source × post-stroke age interaction for the QA/KYNA ratio (F (4, 27) =4.174, p=0.0093). Significant main effects of hemisphere source and post-stroke age were also observed for the KYN/TRYP, QA/KYNA, and KYNA/KYN ratios (**Fig. 3B–D**). *Post hoc* analysis showed that the KYN/TRYP and QA/KYNA ratios were significantly higher in the ipsilateral hemisphere than in the contralateral hemisphere and controls at all time points (p<0.001). Additionally, the KYN/TRYP ratio was significantly elevated in the contralateral hemisphere compared with controls across all post-stroke days (p<0.01), while the QA/KYNA ratio was significantly higher in the contralateral hemisphere than in controls at 1 and 7 days post-stroke (p<0.05). In contrast, the 3-HK/3-HAA ratio did not differ significantly between the ipsilateral hemisphere and controls.

The KYNA/KYN ratio was consistently reduced in the ipsilateral hemisphere at all post-stroke time points relative to the contralateral (p<0.001) and control (p<0.0001) groups **(Fig. 3D)**. Additionally, the KYNA/KYN ratio was lower in the contralateral hemisphere (p<0.0001) when compared with the control across all time points.

For the *in vitro* studies, we first validated the immune-inflammatory state of LPS- and ICM-stimulated BV2 microglia by measuring mRNA expression of key inflammatory genes (**Fig. 4**). One-way ANOVA revealed that LPS and ICM stimulation significantly (p<0.05) upregulated *Nos2, Ptgs2*, *Tnf*, and *Il6* expression compared to PBS (**Fig. 4A**), confirming robust activation of genes associated with an amplifying or pro-inflammatory type response in BV2 microglia. Furthermore, LPS stimulation significantly increased *Il1b* mRNA expression relative to PBS. Consistent with an amplifying response, LPS and ICM treatment, compared with PBS, significantly downregulated *Arg1*, which is associated with a reparative activation profile. (**Fig. 4A**).

**Fig. 4:**
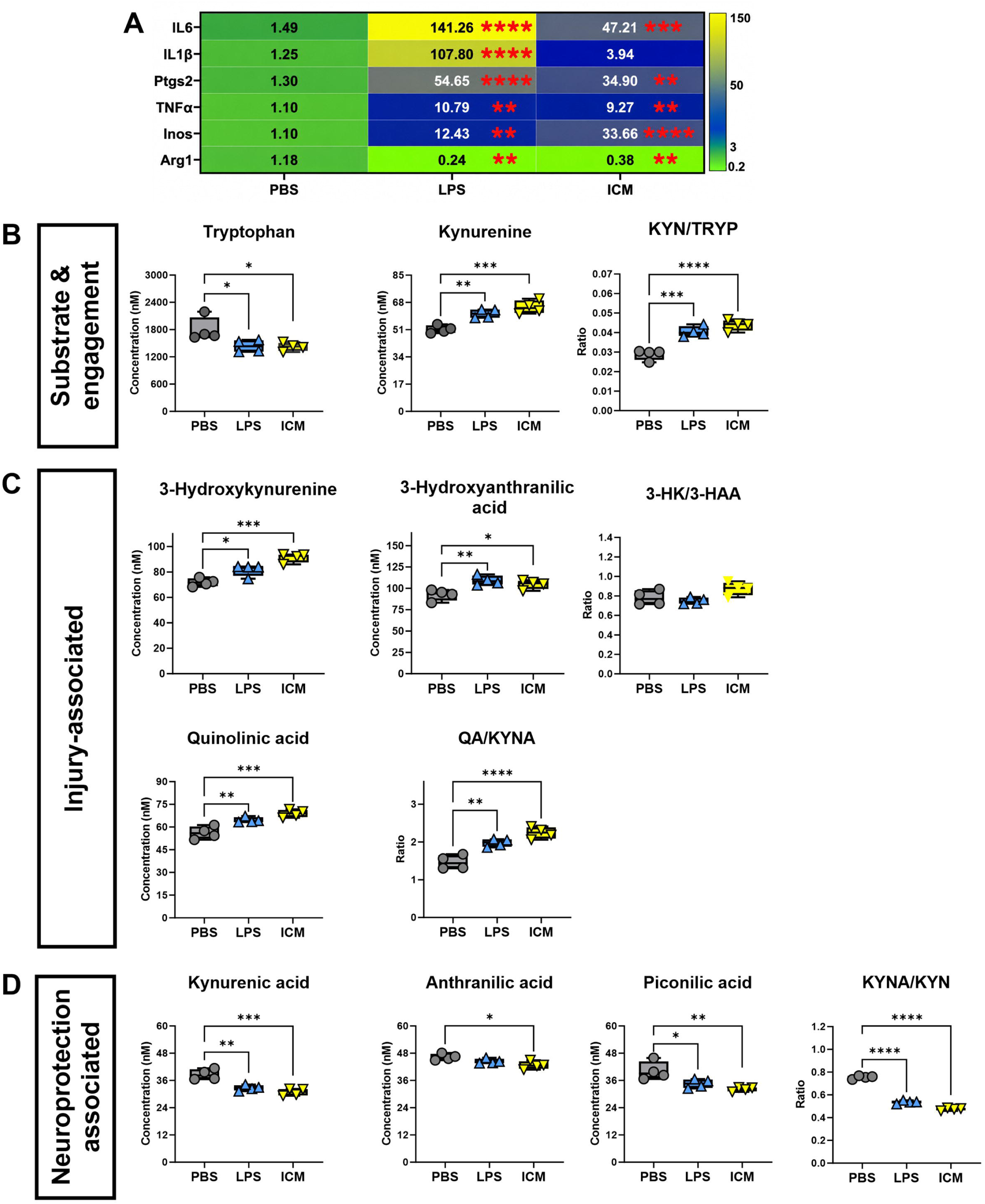
Induction of a pro-inflammatory gene profile in BV2 microglia and quantitative analysis of kynurenine pathway metabolites from the matched cell supernatant. (**A**) Summary of gene expression from BV2 microglia stimulated with lipopolysaccharide (LPS) or a proinflammatory cytokine mix (ICM; IL-1β and IFN-γ). Then in **B**) the KP metabolites associated with substrate and pathway engagement (tryptophan [TRYP], kynurenine [KYN] and the KYN/TRYP ratio), in **C**) metabolites associated with injury progression (3-hydroxykynurenine [3-HK], 3-hydroxyanthranilic acid [3-HAA], the 3-HK/3-HAA ratio, quinolinic acid [QA] and the QA/KYNA ratio) and in **D**) metabolites associated with a neuroprotective response (kynurenic acid [KYNA], anthranilic acid [AA], and picolinic acid [PA], and KYNA/KYN ratio). Data are presented as mean ± SEM; independent experiment n = 4 per group, shown as individual points on the graphs; outlined in Supplementary Table S7 and S8. Statistical significance was determined using a one-way ANOVA followed by Šídák’s multiple comparisons test. Significance in the post-test is shown on the graphs (*p < 0.05, **p < 0.01, ***p < 0.001, ****p <0.0001).

One-way ANOVA demonstrated a significant effect of LPS and ICM on all KP metabolites (p<0.05) (**Fig. 4B**). In Sidak’s post-test, LPS and ICM significantly decreased TRYP levels (p<0.05) and elevated the levels of KYN (p<0.01) and the ratio of KYN/TRYP (p<0.001) in BV2-microglia cell supernatant compared to PBS (**Fig. 4B)**. Similarly, levels of the neurotoxic metabolites 3-HK, 3-HAA, and QA were significantly increased (p<0.05) in supernatants from LPS and ICM-stimulated cells compared with PBS controls. In addition, both LPS and ICM treatments significantly elevated the QA/KYNA ratio (p<0.01) relative to PBS (**Fig. 4C**). In contrast, the LPS and ICM stimulation of BV2-microglia resulted in a significant (p<0.01) reduction in the neuroprotective KYNA and the KYNA/KYN ratio compared with PBS, consistent with the reductions observed in the neonatal stroke brain (**Fig. 4D**). ICM stimulation also resulted in significantly lower AA and PA levels compared with PBS treated controls (p<0.05) (**Fig. 4D).**

## Discussion

Dysregulated KP metabolites have been linked to neurodegenerative diseases, including Alzheimer’s disease ^39^, Parkinson’s disease ^40^, schizophrenia ^41^ and Huntington’s disease ^42^, as well as adverse pregnancy outcomes such as fetal growth restriction and preterm birth ^43^. Emerging evidence also suggests that KP imbalance may contribute to injury progression following early-life brain injury ^44^. Precise quantification of multiple KP metabolites across central nervous system-relevant injury paradigms and biological matrices is therefore essential to improve the measurement and interpretation of KP alterations in health and disease and to support the investigation of metabolic biomarkers.

In this study, we developed and validated a sensitive, accurate, and robust LC-MS/MS method for the integrated quantification of eight KP metabolites and four pathway-balance ratios in two complex biological matrices: brain homogenates from a rat model of neonatal stroke and cell culture supernatants from pro-inflammatory activated microglia. While individual metabolite measurements show specific KP changes, pathway-balance ratios provide additional insight into shifts in molecule entry into the pathway and between the pathway’s neuroprotective, oxidative and neurotoxic branches. The developed LC-MS/MS method shows excellent linearity, sensitivity, imprecision, accuracy, and matrix stability across all eight metabolites, providing a reliable platform for quantitatively assessing KP metabolites in preclinical models of brain injury.

Analysing KP metabolites is challenging because they span a wide polarity range, from hydrophobic TRYP to highly hydrophilic QA, 3-HK, and PA, which complicates chromatographic separation and ionisation ^45^. In this study, we addressed these issues by optimising ionisation conditions and adopting ESI^+^ for all eight KP metabolites. ESI^+^ provided superior fragmentation and sensitivity compared with negative mode, which underperforms for metabolites such as 3-HAA and AA ^46^. Matrix effects in complex samples such as plasma, cerebrospinal fluid, and brain homogenates further complicate KP metabolite analysis by distorting ionisation efficiency and compromising quantitative accuracy^46, 47^. We used a protein precipitation step followed by EMR-lipid Captiva, as this two-step approach improves extraction efficiency and recovery while reducing matrix interference compared with a single preparation method^23, 48, 49^.

Earlier LC–MS/MS methods applied to brain tissue quantified four or five KP metabolites ^50–52^; reported seven KP metabolites, could not quantify 3-HAA, and gave no accuracy, precision or LLOQ data ^53^; or quantified eight KP metabolites, excluding AA, in plasma, CSF and brain while determining accuracy and precision in plasma alone, with a within-run accuracy for 3-HAA of 42–67% ^11^. The only method we identified for cell culture medium quantified four KP metabolites using single-quadrupole LC–MS ^46^, whereas our method quantified all eight KP metabolites, including AA, and we validated it separately in brain homogenate and cell culture supernatant.

These findings demonstrate the method’s sensitivity and applicability across complex biological matrices. In addition to imprecision assessment, our method demonstrated excellent intraday and interday precision, with imprecision consistently <12%, below the CLSI-recommended limit of 15–20% ^52^. Accuracy ranged from 89.2%–108.3% in the brain matrix and 90.84%–95.92% in the cell culture matrix, with recoveries of 83.2–103.1% and 86.66–112.31%, respectively. These findings both meet the CLSI guidelines for accuracy and recovery ^30^, and are consistent with previous studies ^54, 55^.

Stability testing showed that 3-HK, 3-HAA, and QA degraded by more than 15% over 14 days at −20 °C. Based on our data, we recommend that samples for analysis of these compounds be snap-frozen and stored at or below −20 °C (at our testing concentrations) and analysed within 7–14 days ^46, 56, 57^. This variability in stability likely reflects the high intrinsic redox reactivity and low natural abundance of these compounds ^58, 59^.

The method showed strong analytical performance and reproducibility across all validation parameters, supporting its suitability for robust quantification of KP metabolites. As with all targeted LC-MS/MS approaches, interference from structurally related or isobaric compounds cannot be entirely excluded, although chromatographic separation and qualifier ion monitoring were used to minimise this risk. External cross-validation was limited because traceable, commercially available, matrix-matched calibrators, QC, and external quality assurance (EQA) materials for each KP metabolite were unavailable.

To evaluate the method’s applicability, we applied the validated LC-MS/MS method to rat brain tissue homogenates from a neonatal stroke model and BV2-microglia cell culture supernatants from a neuroinflammation model. The method showed enhanced TRYP breakdown, with increased downstream KP metabolites (including KYN, 3-HK, 3-HAA, and QA) and reduced levels of the neuroprotective metabolite KYNA. Although similar KP dysregulation has been described in adult ischemic stroke models ^60, 61^, this is the first use of a single validated LC-MS/MS platform to quantify eight KP metabolites and pathway-balance ratios in neonatal stroke brain tissue.

Consistent with the *in vivo* findings, analysis of supernatant from microglia induced to a pro-inflammatory state revealed a comparable pattern of KP metabolite changes, characterised by increased production of neurotoxic KP metabolites and reduced levels of the neuroprotective metabolite KYNA. Our findings align with previous studies showing that LPS stimulation of BV2 or primary mouse microglia significantly increased KYN and QA ^62^. We tested the effects of LPS and combined IL-1β and IFN-γ on KP metabolites as models of bacterial sepsis and a moderate sterile inflammatory response, respectively. Both these paradigms are commonly used to model aspects of neuroinflammation in perinatal and adult brain insults ^36, 63–65^. Although previous studies have shown changes in KP following immune activation in microglia, most were limited to a single immune stimulus or a small number of metabolites ^62^. By enabling quantification of multiple KP metabolites, our results show this analytical approach can capture pathway-level changes that mirror those observed *in vivo* following neonatal brain injury. Collectively, these findings highlight this analytical platform’s utility for characterising KP alterations across experimental models of brain injury, positioning it as a valuable tool for future mechanistic exploration and translational research.

## Conclusion

In summary, we present a rigorously validated LC-MS/MS assay that enables sensitive, accurate, and reproducible quantification of key KP metabolites across complex biological matrices. This method robustly quantifies eight metabolites in neonatal stroke brain tissue and immune-activated BV2-microglia supernatants, making it a valuable tool for assessing KP metabolism in neurological disorders. Application of the method identified patterns consistent with a shift toward a neurotoxic KP profile in both *in vivo* and *in vitro* contexts, supporting its suitability for investigating metabolic changes associated with brain injury. Together, this platform offers a valuable analytical tool for investigating KP dynamics in neuroinflammatory, neurodevelopmental, and neurodegenerative disorders.

## Supporting information

Supplementary_Figures_and_Tables_Hossen_Fleiss_Zakaria_biorxiv_2026

## Acknowledgements

The authors thank all collaborators and the technical staff of the School of Health and Biomedical Sciences, RMIT University, for their support for this work. During the preparation of this manuscript, the authors used Val (RMIT University’s internal generative AI tool, built on OpenAI’s GPT-5.0 model) and Grammarly (1.205.1.0, WebUI 2.46.0) to check and correct the grammar and spelling. The tool was not used to generate content, analyse data, or interpret results. The authors reviewed and edited all output and take full responsibility for the content of the publication.

## Author contributions

MMH: Conceptualisation, methodology, formal analysis, data curation, validation, visualisation, investigation, and writing the original draft. VM, NE, RK, ERM, AAMS, and IKS: Methodology, analysis, data curation, and manuscript review and editing. BF and RZ: Conceptualisation, supervision, funding acquisition, data curation, formal analysis, validation, visualisation, project administration, and manuscript writing, review and editing. All authors reviewed and approved the final manuscript.

## Statements and Declarations

### Ethical considerations

All animal procedures were approved by the RMIT University Animal Ethics Committee (approval number AEC 27678) and are reported in accordance with the ARRIVE guidelines. No human participants, human data or human tissue were used.

### Consent to participate

Not applicable.

### Consent for publication

Not applicable.

### Data availability

The data supporting the findings of this study are available from the corresponding authors on reasonable request.

