## Supplementary_Figures_and_Tables_Hossen_Fleiss_Zakaria_biorxiv_2026 for "Validated LC-MS/MS quantification of kynurenine pathway metabolites and pathway-balance ratios in brain tissue and microglial culture supernatant: application to neonatal stroke"

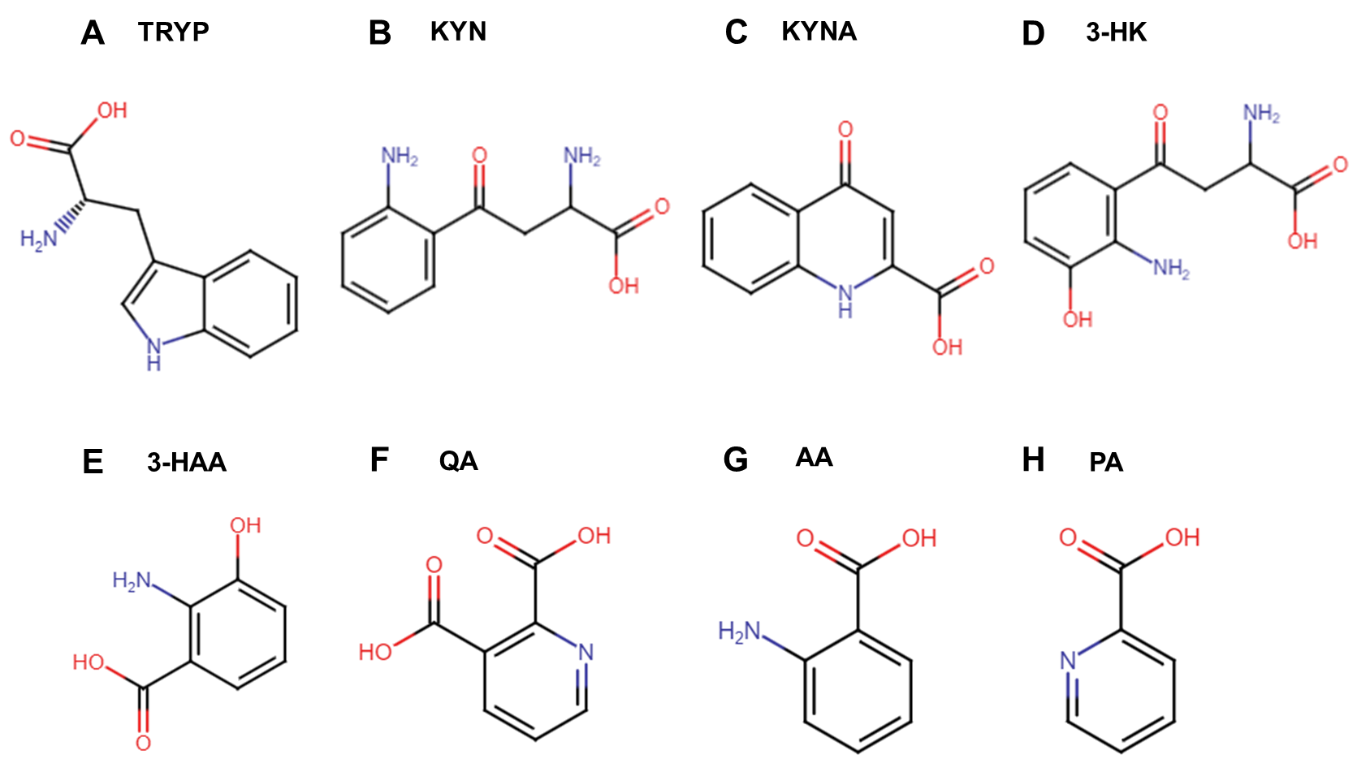

**Supplementary Figure S1:** Chemical structure of kynurenine pathway metabolites. (A) TRYP, tryptophan; (B) KYN, kynurenine; (C) KYNA, kynurenic acid; (D) 3-HK, 3-hydroxykynurenine; (E) 3-HAA, 3-hydroxyanthranilic acid; (F) QA, quinolinic acid; (G) AA, anthranilic acid; and (H) PA, picolinic acid. Nitrogen atoms are shown in blue and oxygen atoms in red; carbon atoms are shown in black.

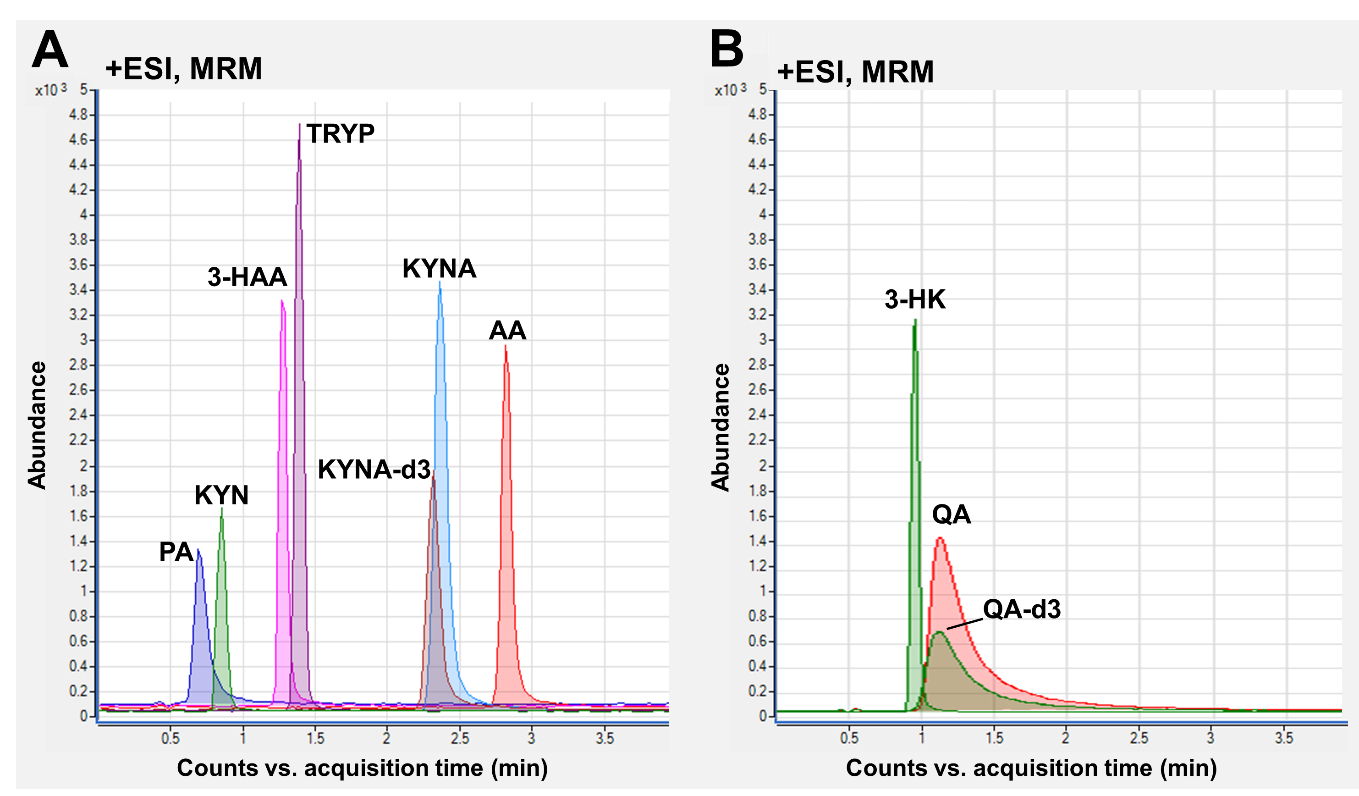

**Supplementary Figure S2:** Representative LC–MS/MS chromatograms obtained using two different mobile phase gradients. (A) Chromatographic separation achieved using mobile phase gradient 1. (B) Chromatographic separation achieved using mobile phase gradient 2. Peaks correspond to targeted KP metabolites, identified based on retention time and specific MRM transitions. Tryptophan (TRYP), kynurenine (KYN), kynurenic acid (KYNA), Quinolinic acid (QA), 3-hydroxykynurenine (3-HK), 3-hydroxyanthranilic acid (3-HAA), anthranilic acid (AA), picolinic acid (PA), Electrospray ionization (ESI), and Multiple reaction monitoring (MRM).

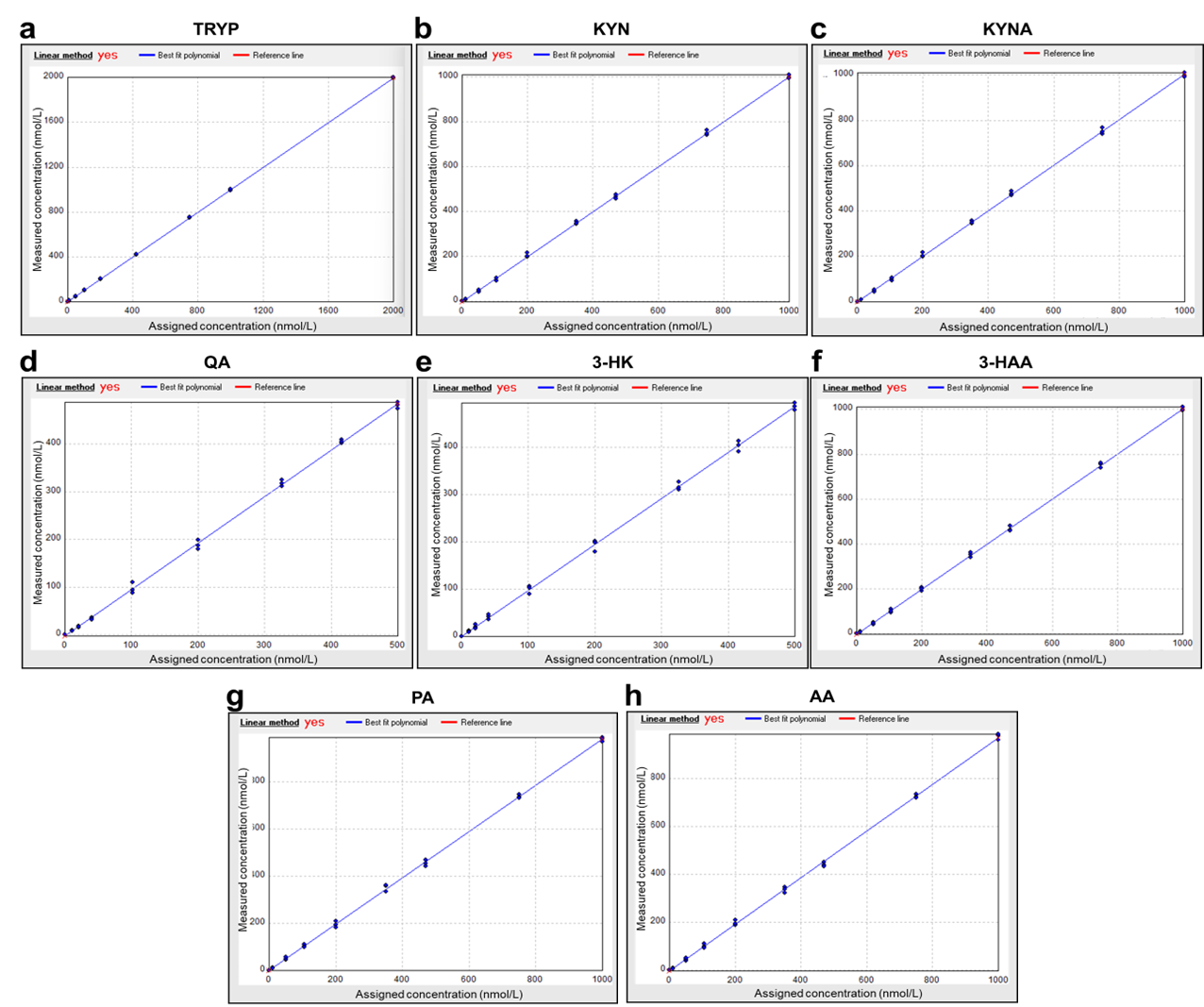

**Supplementary Figure S3:** Representative linear curves for kynurenine pathway metabolites. Calibration curves for a) tryptophan (TRYP), b) kynurenine (KYN), c) kynurenic acid (KYNA), d) quinolinic acid (QA), e) 3-hydroxykynurenine (3-HK), f) 3-hydroxyanthranilic acid (3-HAA), g) picolinic acid (PA), and h) anthranilic acid (AA).

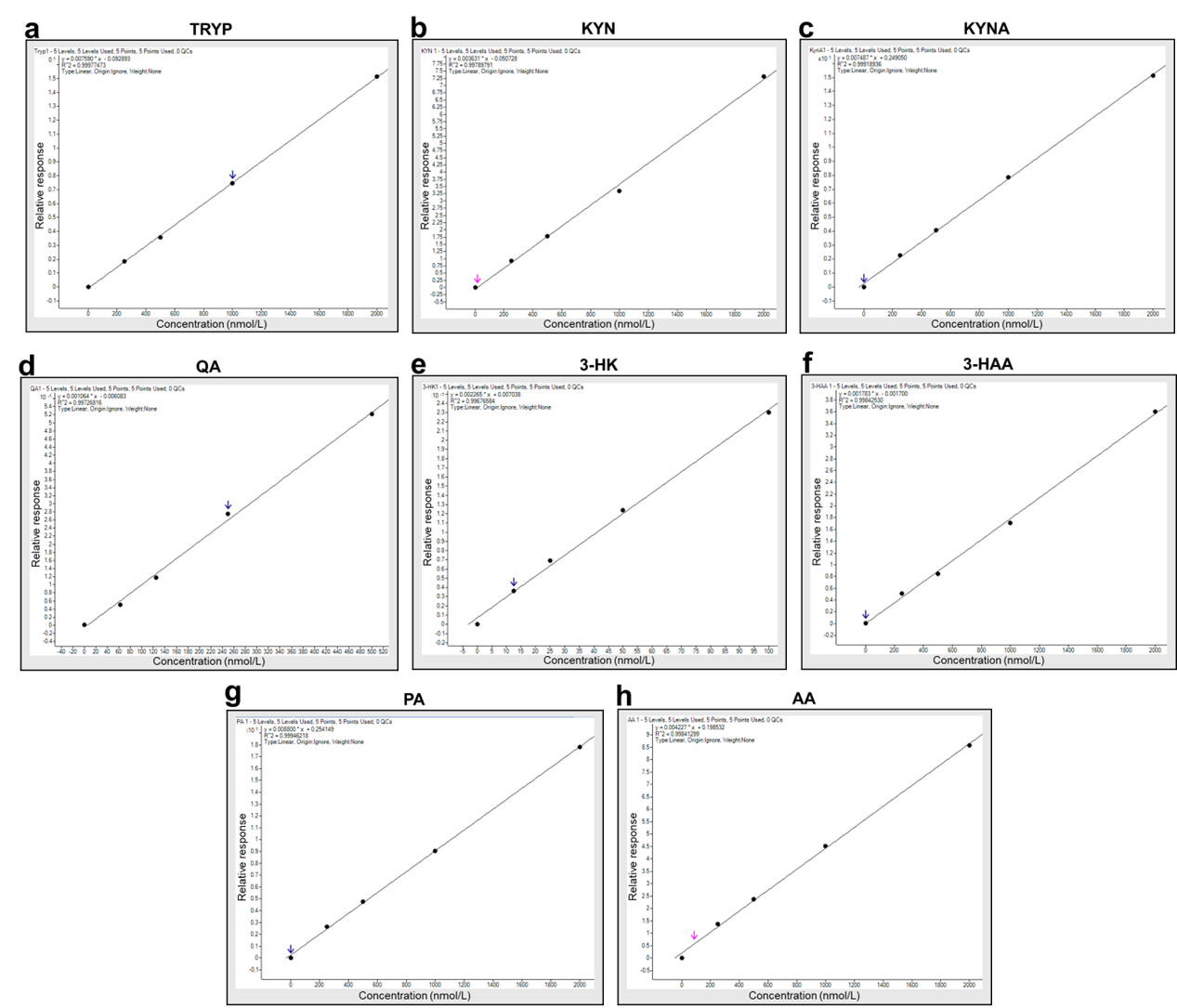

**Supplementary Figure S4:** Representative calibration curves for kynurenine pathway metabolites. Calibration curves for a) tryptophan (TRYP), b) kynurenine (KYN), c) kynurenic acid (KYNA), d) quinolinic acid (QA), e) 3-hydroxykynurenine (3-HK), f) 3-hydroxyanthranilic acid (3-HAA), g) picolinic acid (PA), and h) anthranilic acid (AA).

| **Supplementary Table S1**: Summary of kynurenine pathway metabolites, standards and solvents/diluents | | | | | |
| --- | --- | --- | --- | --- | --- |
| **Metabolite** | **Molecular Mass (g)** | **Solvent for STD diluent** | **Volume (mL)** | **1st Stock Conc. (nM)** | **2nd Stock Conc. (nM)** |
| TRYP | 204.23 | Water: Methanol (50:50) +0.1% FA | 50 | 979288 | 20000 |
| KYN | 208.21 | Water: Methanol (50:50) +0.1% FA | 25 | 960568 | 20000 |
| KYNA | 189.17 | Water | 50 | 105725 | 20000 |
| KYNA-d3 | 194.20 | Methanol | 25 | 514933 | 20000 |
| QA | 167.12 | Methanol | 100 | 30690521 | 20000 |
| QA-d3 | 170.14 | Methanol | 25 | 235100 | 20000 |
| 3-HK | 224.21 | Water: Methanol (80:20) + 0.1% FA | 25 | 892020 | 20000 |
| PA | 123.11 | Methanol | 100 | 40037364 | 20000 |
| 3-HAA | 153.14 | Methanol | 100 | 6529972 | 20000 |
| AA | 137.14 | Water: Methanol (70:30) + 0.1% FA | 50 | 1458363 | 20000 |
| **Abbreviations:** Tryptophan (TRYP), kynurenine (KYN), kynurenic acid (KYNA), Quinolinic acid (QA), 3-hydroxykynurenine (3-HK), 3-hydroxyanthranilic acid (3-HAA), anthranilic acid (AA), and picolinic acid (PA), Conc., concentration. | | | | | |

| **Supplementary Table S2** Performance specifications of Agilent LC-1290 | | |
| --- | --- | --- |
| **Specification*** | | **Agilent LC-1290 (309-311)** |
| Pump | Hydraulic system | Two dual pistons in series, pumps with proprietary servo-controlled variable stroke design and smooth motion control |
|  | Settable flow range | 0.001 – 5 mL/min, in 0.001 mL/min increments |
|  | Flow precision | ≤0.07 % RSD |
|  | Flow accuracy | ±1% |
|  | Pressure operating range | Binary Pump: up to 120 MPa (1200 bar) up to 2 mL/min,  80 MPa (800 bar) at 5 mL/min  Binary Pump VL: up to 105 MPa (1050 bar) up to 2 mL/min |
|  | Pressure pulsation | < 1 % amplitude |
|  | Gradient formation | High-pressure binary mixing |
|  | Delay volume | JetWeaver V35: < 45μL, JetWeaver V100: < 75μL |
|  | Composition range | Settable range: 0 - 100 % |
|  | Composition precision | < 0.15 % RSD or 0.01 min SD |
| Auto-sampler | Injection range | 0.1–20 μL in 0.1 μL increments |
|  | Injection precision | <0.3%, 5–40μL; <0.5%, 2–5μL; <0.7%, 0.5–2μL |
|  | Sample viscosity range | 0.2–5 centipoise |
|  | Injection cycle time | Typically, < 21 s depending on draw speed and injection vol. |
|  | Carryover | Typically, < 0.004 % |
| Column Oven | Temperature range | 10 °C below ambient to 100°C |
|  | Temperature stability | ± 0.05°C |
|  | Temperature accuracy | ± 0.8°C |
|  | Column capacity | 2 columns of 300 mm with individual solvent heating |
|  | Internal volume | 3 μL left heat exchanger, 6 μL right heat exchanger |

* These performance specifications are based on data analysed by Agilent Technologies.

| **Supplementary Table S3**: Performance specifications of Agilent MS/MS QQQ-6490. | |
| --- | --- |
| **Specification*** | **Agilent QQQ-6490 (312)** |
| Sensitivity: 1pg reserpine# quantifying on m/z 609→195 using ESI positive mode | S/N > 50,000:1 |
| Sensitivity: 1pg chloramphenicol# quantifying on m/z 321→152 using ESI negative mode | S/N > 50,000:1 |
| Linear dynamic range | > 6.0 × 10^6^ |
| Mass range | 5–1,400 Da |
| Maximum scan rate | 12,500 Da/sec |
| Minimum MRM dwell time | 1 millisecond |
| MRM transitions | 500 per TS, >40,000 ion transitions per method |
| Polarity switching (from positive to negative) | 30 milliseconds |
| Collision cell ion clearance | < 1 millisecond |
| Agilent Jet stream technology* | Yes |
| Agilent iFunnel technology* | Yes |
| * Agilent Technologies uses reserpine and chloramphenicol compounds to compare their MS/MS systems. | |

| **Supplementary Table S4:** Results of the statistical analyses for tryptophan, kynurenine, and the KYN/TRP ratio across control, contralateral, and ipsilateral hemispheres. | | | | | | | | | |
| --- | --- | --- | --- | --- | --- | --- | --- | --- | --- |
| **Figure** | **KP metabolites** | **Test name** | **Main effect** | | | **Sidak’s multiple comparison** | | | |
|  |  |  | **Interaction, F (DFn, DFd),**  **p value** | **Hemisphere, F (DFn, DFd),**  **p value** | **Time**  **F (DFn,**  **DFd), p value** | **Post-stroke time** | **Hemisphere sources** | **95% CI** | **p value** |
| **Figure 3B,** Substrate & engagement | Tryptophan | Mixed-effect analysis | F (4, 6) = 1.855,  p = 0.2376 | F (2, 6) = 98.34,  *p < 0.0001* | F (2, 21) = 55.56,  *p < 0.0001* | 1 day | Control vs. Contralateral | 192.2 to 607.2 | *0.0022* |
|  |  |  |  |  |  |  | Control vs. Ipsilateral | 376.0 to 791.0 | *0.0003* |
|  |  |  |  |  |  |  | Contralateral vs. Ipsilateral | 84.57 to 283.1 | *0.0027* |
|  |  |  |  |  |  | 7 days | Control vs. Contralateral | 78.14 to 493.1 | *0.0122* |
|  |  |  |  |  |  |  | Control vs. Ipsilateral | 228.8 to 643.7 | *0.0014* |
|  |  |  |  |  |  |  | Contralateral vs. Ipsilateral | 51.35 to 249.9 | *0.0076* |
|  |  |  |  |  |  | 14 days | Control vs. Contralateral | -16.25 to 398.7 | *0.0688* |
|  |  |  |  |  |  |  | Control vs. Ipsilateral | 136.9 to 551.8 | *0.0048* |
|  |  |  |  |  |  |  | Contralateral vs. Ipsilateral | 53.87 to 252.4 | *0.0070* |
|  | Kynurenine | Mixed-effect analysis | F (4, 6) = 0.6775,  p = 0.6320 | F (2, 6) = 169.0,  *p < 0.001* | F (2, 21) = 4.072,  *p < 0.0320* | 1 day | Control vs. Contralateral | -14.59 to 2.701 | *0.1835* |
|  |  |  |  |  |  |  | Control vs. Ipsilateral | -24.34 to -7.046 | 0.0030 |
|  |  |  |  |  |  |  | Contralateral vs. Ipsilateral | -13.16 to -6.338 | 0.0003 |
|  |  |  |  |  |  | 7 days | Control vs. Contralateral | -15.38 to 1.910 | *0.1248* |
|  |  |  |  |  |  |  | Control vs. Ipsilateral | -26.23 to -8.940 | 0.0017 |
|  |  |  |  |  |  |  | Contralateral vs. Ipsilateral | -14.26 to -7.441 | 0.0001 |
|  |  |  |  |  |  | 14 days | Control vs. Contralateral | -18.64 to -1.349 | 0.0271 |
|  |  |  |  |  |  |  | Control vs. Ipsilateral | -27.58 to -10.29 | 0.0011 |
|  |  |  |  |  |  |  | Contralateral vs. Ipsilateral | -12.35 to -5.531 | 0.0004 |
|  | KYN/TRYP | Mixed-effect analysis | F (4, 6) = 1.300,  p = 0.3680 | F (2, 6) = 291.4,  *p <0.0001* | F (2, 21) = 11.77,  *p < 0.0004* | 1 day | Control vs. Contralateral | -0.011 to -0.002 | 0.0075 |
|  |  |  |  |  |  |  | Control vs. Ipsilateral | -0.02 to -0.011 | 0.0001 |
|  |  |  |  |  |  |  | Contralateral vs. Ipsilateral | -0.011 to -0.005 | 0.0004 |
|  |  |  |  |  |  | 7 days | Control vs. Contralateral | -0.011 to -0.003 | 0.0049 |
|  |  |  |  |  |  |  | Control vs. Ipsilateral | -0.022 to -0.013 | 0.0001 |
|  |  |  |  |  |  |  | Contralateral vs. Ipsilateral | -0.013 to -0.007 | 0.0001 |
|  |  |  |  |  |  | 14 days | Control vs. Contralateral | -0.014 to -0.004 | 0.0017 |
|  |  |  |  |  |  |  | Control vs. Ipsilateral | -0.024 to -0.015 | 0.0001 |
|  |  |  |  |  |  |  | Contralateral vs. Ipsilateral | -0.013 to -0.007 | 0.0001 |

| **Supplementary Table S5:** Results of the statistical analyses for injury associated KP metabolites and ratios across control, contralateral, and ipsilateral hemispheres. | | | | | | | | | |
| --- | --- | --- | --- | --- | --- | --- | --- | --- | --- |
| **Figure** | **KP metabolites** | **Test name** | **Main effect** | | | **Sidak’s multiple comparison** | | | |
|  |  |  | **Interaction, F (DFn, DFd),**  **p value** | **Hemisphere, F (DFn, DFd),**  **p value** | **Time**  **F (DFn,**  **DFd), p value** | **Post-stroke time** | **Hemisphere sources** | **95% CI** | **p value** |
| **Figure 3C,** Injury associated | 3-Hydroxykynurenine | Mixed-effect analysis | F (4, 6) = 0.9409,  p = 0.5004 | F (2, 6) = 64.41,  *p < 0.0001* | F (2, 21) = 23.38,  *p < 0.0001* | 1 day | Control vs. Contralateral | -10.61 to 0.89 | *0.0946* |
|  |  |  |  |  |  |  | Control vs. Ipsilateral | -14.77 to -3.26 | *0.0064* |
|  |  |  |  |  |  |  | Contralateral vs. Ipsilateral | -7.66 to -0.65 | *0.0242* |
|  |  |  |  |  |  | 7 days | Control vs. Contralateral | -12.46 to -0.95 | *0.0260* |
|  |  |  |  |  |  |  | Control vs. Ipsilateral | -17.22 to -5.71 | *0.0019* |
|  |  |  |  |  |  |  | Contralateral vs. Ipsilateral | -8.25 to -1.24 | *0.0131* |
|  |  |  |  |  |  | 14 days | Control vs. Contralateral | -9.04 to 2.45 | *0.2947* |
|  |  |  |  |  |  |  | Control vs. Ipsilateral | -15.47 to -3.96 | *0.0044* |
|  |  |  |  |  |  |  | Contralateral vs. Ipsilateral | -9.92 to -2.91 | *0.0029* |
|  | 3-Hydroxyanthranilic acid | Mixed-effect analysis | F (4, 6) = 0.5031,  p = 0.736 | F (2, 6) = 77.44,  *p < 0.0001* | F (2, 21) = 5.226,  *p < 0.014* | 1 day | Control vs. Contralateral | -8.393 to 2.113 | *0.2665* |
|  |  |  |  |  |  |  | Control vs. Ipsilateral | -13.87 to -3.36 | *0.0051* |
|  |  |  |  |  |  |  | Contralateral vs. Ipsilateral | -8.11 to -2.84 | *0.0015* |
|  |  |  |  |  |  | 7 days | Control vs. Contralateral | -10.49 to 0.01 | *0.0505* |
|  |  |  |  |  |  |  | Control vs. Ipsilateral | -14.73 to -4.22 | *0.0031* |
|  |  |  |  |  |  |  | Contralateral vs. Ipsilateral | -6.87 to -1.60 | *0.0057* |
|  |  |  |  |  |  | 14 days | Control vs. Contralateral | -8.42 to 2.08 | *0.2596* |
|  |  |  |  |  |  |  | Control vs. Ipsilateral | -13.09 to -2.58 | *0.0082* |
|  |  |  |  |  |  |  | Contralateral vs. Ipsilateral | -7.30 to -2.03 | *0.0035* |
|  | 3-HK/3-HAA | Mixed-effect analysis | F (4, 6) = 2.007*,*  p = 0.212 | F (2, 6) = 3.663,  *p < 0.091* | F (2, 21) = 5.474,  *p < 0.012* | 1 day | Control vs. Contralateral | -0.38 to 0.16 | *0.5566* |
|  |  |  |  |  |  |  | Control vs. Ipsilateral | -0.35 to 0.19 | *0.7676* |
|  |  |  |  |  |  |  | Contralateral vs. Ipsilateral | -0.068 to 0.13 | *0.7154* |
|  |  |  |  |  |  | 7 days | Control vs. Contralateral | -0.38 to 0.16 | *0.5485* |
|  |  |  |  |  |  |  | Control vs. Ipsilateral | -0.41 to 0.12 | *0.3546* |
|  |  |  |  |  |  |  | Contralateral vs. Ipsilateral | -0.13 to 0.066 | *0.6849* |
|  |  |  |  |  |  | 14 days | Control vs. Contralateral | -0.30 to 0.24 | *0.9773* |
|  |  |  |  |  |  |  | Control vs. Ipsilateral | -0.39 to 0.15 | *0.4852* |
|  |  |  |  |  |  |  | Contralateral vs. Ipsilateral | -0.18 to 0.01 | *0.0776* |
| **Figure 3C,** Injury associated | Quinolinic acid | Mixed-effect analysis | F (4, 6) = 2.481,  p = 0.1535 | F (2, 6) = 152.4,  *p < 0.0001* | F (2, 21) = 32.96,  *p < 0.0001* | 1 day | Control vs. Contralateral | -16.32 to -2.98 | *0.0095* |
|  |  |  |  |  |  |  | Control vs. Ipsilateral | -28.75 to -15.4 | *0.0001* |
|  |  |  |  |  |  |  | Contralateral vs. Ipsilateral | -18.91 to -5.93 | *0.0023* |
|  |  |  |  |  |  | 7 days | Control vs. Contralateral | -19.74 to -6.40 | *0.0020* |
|  |  |  |  |  |  |  | Control vs. Ipsilateral | -30.40 to -17.0 | *0.0001* |
|  |  |  |  |  |  |  | Contralateral vs. Ipsilateral | -17.15 to -4.18 | *0.0051* |
|  |  |  |  |  |  | 14 days | Control vs. Contralateral | -13.46 to -0.12 | *0.0463* |
|  |  |  |  |  |  |  | Control vs. Ipsilateral | -22.35 to -9.01 | *0.0008* |
|  |  |  |  |  |  |  | Contralateral vs. Ipsilateral | -15.37 to -2.40 | *0.0125* |
|  | QA/KYNA | Mixed-effect analysis | F (4, 27) = 4.174,  *p = 0.0092* | F (2, 27) = 121.4,  *p < 0.0001* | F (2, 27) = 2.221,  p = 0.1279 | 1 day | Control vs. Contralateral | -0.89 to -0.04 | *0.0275* |
|  |  |  |  |  |  |  | Control vs. Ipsilateral | -1.95 to -1.10 | *0.0001* |
|  |  |  |  |  |  |  | Contralateral vs. Ipsilateral | -1.49 to -0.64 | *0.0001* |
|  |  |  |  |  |  | 7 days | Control vs. Contralateral | -0.92 to -0.07 | *0.0170* |
|  |  |  |  |  |  |  | Control vs. Ipsilateral | -2.29 to -1.44 | *0.0001* |
|  |  |  |  |  |  |  | Contralateral vs. Ipsilateral | -1.79 to -0.94 | *0.0001* |
|  |  |  |  |  |  | 14 days | Control vs. Contralateral | -0.77 to 0.07 | ***0.1250*** |
|  |  |  |  |  |  |  | Control vs. Ipsilateral | -1.40 to -0.55 | *0.0001* |
|  |  |  |  |  |  |  | Contralateral vs. Ipsilateral | -1.05 to -0.20 | *0.0024* |

| **Supplementary Table S6:** Results of the statistical analyses for kynurenic acid and KYNA/KYN ratio across control, contralateral, and ipsilateral hemispheres. | | | | | | | | | |
| --- | --- | --- | --- | --- | --- | --- | --- | --- | --- |
| **Figure** | **KP metabolites** | **Test name** | **Main effect** | | | **Sidak’s multiple comparison** | | | |
|  |  |  | **Interaction, F (DFn, DFd),**  **p value** | **Hemisphere, F (DFn, DFd),**  **p value** | **Time**  **F (DFn,**  **DFd), p value** | **Post-stroke time** | **Hemisphere sources** | **95% CI** | **p value** |
| **Figure 3D,**  Neuroprotection associated | Kynurenic acid | Mixed-effect analysis | F (4, 6) = 3.512,  p = 0.066 | F (2, 6) = 162.1,  *p < 0.0001* | F (2, 21) = 17.68,  *p < 0.0001* | 1 day | Control vs. Contralateral | 4.95 to 18.33 | 0.0038 |
|  |  |  |  |  |  |  | Control vs. Ipsilateral | 18.67 to 32.05 | 0.0001 |
|  |  |  |  |  |  |  | Contralateral vs. Ipsilateral | 7.16 to 20.27 | 0.0014 |
|  |  |  |  |  |  | 7 days | Control vs. Contralateral | 2.36 to 15.73 | 0.0132 |
|  |  |  |  |  |  |  | Control vs. Ipsilateral | 18.90 to 32.28 | 0.0001 |
|  |  |  |  |  |  |  | Contralateral vs. Ipsilateral | 9.98 to 23.09 | 0.0005 |
|  |  |  |  |  |  | 14 days | Control vs. Contralateral | -1.08 to 12.29 | 0.0977 |
|  |  |  |  |  |  |  | Control vs. Ipsilateral | 5.58 to 18.96 | 0.0029 |
|  |  |  |  |  |  |  | Contralateral vs. Ipsilateral | 0.12 to 13.23 | 0.0463 |
|  | KYNA/KYN | Mixed-effect analysis | F (4, 27) = 2.189,  p = 0.0971 | F (2, 27) = 227.1,  *p < 0.0001* | F (2, 27) = 1.357,  p = 0.2744 | 1 day | Control vs. Contralateral | 0.20 to 0.48 | *0.0001* |
|  |  |  |  |  |  |  | Control vs. Ipsilateral | 0.56 to 0.85 | *0.0001* |
|  |  |  |  |  |  |  | Contralateral vs. Ipsilateral | 0.22 to 0.50 | *0.0001* |
|  |  |  |  |  |  | 7 days | Control vs. Contralateral | 0.20 to 0.49 | *0.0001* |
|  |  |  |  |  |  |  | Control vs. Ipsilateral | 0.64 to 0.92 | *0.0001* |
|  |  |  |  |  |  |  | Contralateral vs. Ipsilateral | 0.29 to 0.57 | *0.0001* |
|  |  |  |  |  |  | 14 days | Control vs. Contralateral | 0.20 to 0.49 | *0.0001* |
|  |  |  |  |  |  |  | Control vs. Ipsilateral | 0.43 to 0.72 | *0.0001* |
|  |  |  |  |  |  |  | Contralateral vs. Ipsilateral | 0.09 to 0.37 | *0.0008* |
|  | AA | Mixed-effect analysis | F (4, 6) = 2.009,  p = 0.2123 | F (2, 6) = 0.7380,  p = 0.5170 | F (2, 21) = 1.170,  p = 0.3299 | 1 day | Control vs. Contralateral | -3.11 to 3.48 | 0.9973 |
|  |  |  |  |  |  |  | Control vs. Ipsilateral | -3.87 to 2.71 | 0.9286 |
|  |  |  |  |  |  |  | Contralateral vs. Ipsilateral | -2.17 to 0.64 | 0.3320 |
|  |  |  |  |  |  | 7 days | Control vs. Contralateral | -1.99 to 4.59 | 0.5696 |
|  |  |  |  |  |  |  | Control vs. Ipsilateral | -2.32 to 4.26 | 0.7525 |
|  |  |  |  |  |  |  | Contralateral vs. Ipsilateral | -1.73 to 1.08 | 0.8567 |
|  |  |  |  |  |  | 14 days | Control vs. Contralateral | -2.67 to 3.91 | 0.9153 |
|  |  |  |  |  |  |  | Control vs. Ipsilateral | -1.88 to 4.70 | 0.5104 |
|  |  |  |  |  |  |  | Contralateral vs. Ipsilateral | -0.62 to 2.19 | 0.3119 |
|  | PA | Mixed-effect analysis | F (4, 6) = 0.6110,  p = 0.6703 | F (2, 6) = 1.060,  p = 0.4036 | F (2, 21) = 3.811,  p = 0.0587 | 1 day | Control vs. Contralateral | -3.34 to 6.28 | 0.7331 |
|  |  |  |  |  |  |  | Control vs. Ipsilateral | -4.46 to 5.17 | 0.9939 |
|  |  |  |  |  |  |  | Contralateral vs. Ipsilateral | -3.79 to 1.56 | 0.5300 |
|  |  |  |  |  |  | 7 days | Control vs. Contralateral | -4.10 to 5.52 | 0.9564 |
|  |  |  |  |  |  |  | Control vs. Ipsilateral | -4.66 to 4.96 | 0.9996 |
|  |  |  |  |  |  |  | Contralateral vs. Ipsilateral | -3.24 to 2.12 | 0.8892 |
|  |  |  |  |  |  | 14 days | Control vs. Contralateral | -6.37 to 3.26 | 0.7008 |
|  |  |  |  |  |  |  | Control vs. Ipsilateral | -6.74 to 2.88 | 0.5557 |
|  |  |  |  |  |  |  | Contralateral vs. Ipsilateral | -3.06 to 2.30 | 0.9611 |

| **Supplementary Table S7:** Results of the statistical analyses for inflammatory and anti-inflammatory genes in PBS, LPS, and ICM treated group. | | | | | |
| --- | --- | --- | --- | --- | --- |
| **Figure** | **Test name** | **Gene** | **Sidak’s multiple comparison** | | |
|  |  |  | **Treatment** | **95% CI** | **p value** |
| **Figure 4A,**  Inflammatory and anti-inflammatory genes | Ordinary one-way ANOVA | IL6 | PBS vs. LPS | -159.1 to -120.5 | <0.0001 |
|  |  |  | PBS vs. ICM | -65.01 to -26.42 | 0.0003 |
|  |  | IL1β | PBS vs. LPS | -125.8 to -87.34 | <0.0001 |
|  |  |  | PBS vs. ICM | -21.91 to 16.52 | 0.9193 |
|  |  | Ptgs2 | PBS vs. LPS | -73.02 to -33.69 | <0.0001 |
|  |  |  | PBS vs. ICM | -53.27 to -13.93 | 0.0027 |
|  |  | TNFα | PBS vs. LPS | -14.62 to -4.77 | 0.0010 |
|  |  |  | PBS vs. ICM | -13.09 to -3.25 | 0.0032 |
|  |  | Inos | PBS vs. LPS | -19.76 to -2.90 | 0.0115 |
|  |  |  | PBS vs. ICM | -41.00 to -24.14 | <0.0001 |
|  |  | Arg1 | PBS vs. LPS | 0.441 to 1.44 | 0.0014 |
|  |  |  | PBS vs. ICM | 0.308 to 1.31 | 0.0038 |

| **Supplementary Table S8:** Results of the statistical analyses for the kynurenine pathway metabolites in PBS, LPS, and ICM treated group. | | | | | |
| --- | --- | --- | --- | --- | --- |
| **Figure** | **Test name** | **Metabolites** | **Sidak’s multiple comparison** | | |
|  |  |  | **Treatment** | **95% CI** | **p value** |
| **Figure 4B,**  Substrate & engagement | Ordinary one-way ANOVA | Tryptophan | PBS vs. LPS | 17.25 to 701.3 | 0.0402 |
|  |  |  | PBS vs. ICM | 33.16 to 717.2 | 0.0329 |
|  |  | Kynurenine | PBS vs. LPS | -15.84 to -3.78 | 0.0037 |
|  |  |  | PBS vs. ICM | -19.89 to -7.83 | 0.0003 |
|  |  | KYN/TRYP | PBS vs. LPS | -0.016 to -0.006 | 0.0004 |
|  |  |  | PBS vs. ICM | -0.02 to -0.009 | <0.0001 |
| **Figure 4C,**  Injury associated | Ordinary one-way ANOVA | 3-Hydroxykynurenine | PBS vs. LPS | -17.22 to -2.58 | 0.0111 |
|  |  |  | PBS vs. ICM | -26.78 to -12.14 | 0.0001 |
|  |  | 3-Hydroxyanthralinic acid | PBS vs. LPS | -27.64 to -5.92 | 0.0050 |
|  |  |  | PBS vs. ICM | -23.11 to -1.39 | 0.0287 |
|  |  | 3-HK/3-HAA | PBS vs. LPS | -0.08 to 0.14 | 0.7316 |
|  |  |  | PBS vs. ICM | -0.21 to 0.022 | 0.1147 |
|  |  | Quinolinic acid | PBS vs. LPS | -14.03 to -2.72 | 0.0065 |
|  |  |  | PBS vs. ICM | -18.42 to -7.12 | 0.0004 |
|  |  | QA/KYNA | PBS vs. LPS | -0.79 to -0.22 | 0.0020 |
|  |  |  | PBS vs. ICM | -1.06 to -0.49 | <0.0001 |
| **Figure 4D,**  Neuroprotection associated | Ordinary one-way ANOVA | Kynurenic acid | PBS vs. LPS | 2.54 to 9.22 | 0.0022 |
|  |  |  | PBS vs. ICM | 4.39 to 11.07 | 0.0003 |
|  |  | Anthranilic acid | PBS vs. LPS | -0.77 to 4.56 | 0.1724 |
|  |  |  | PBS vs. ICM | 0.92 to 6.27 | 0.0114 |
|  |  | Picolinic acid | PBS vs. LPS | 0.60 to 10.57 | 0.0296 |
|  |  |  | PBS vs. ICM | 3.02 to 12.99 | 0.0040 |
|  |  | KYNA/KYN | PBS vs. LPS | 0.19 to 0.24 | <0.0001 |
|  |  |  | PBS vs. ICM | 0.25 to 0.30 | <0.0001 |
